# Defining the importance of the arginine loop region of protegrin-1 for antimicrobial activity towards colistin-resistant *Klebsiella pneumoniae*

**DOI:** 10.1101/2025.05.28.656599

**Authors:** Christina DeBarro, Sofia Assent, Hannah Makhecha, Noor Radde, Rakesh Krishnan, Justin Randall, Cesar de la Fuente-Nunez, Renee M. Fleeman

## Abstract

*Carbapenem resistant Klebsiella pneumoniae* extreme drug resistance has warranted the use of colistin as a last-resort antibiotic but these isolates are now displaying increasing rates of colistin resistance. This is in large part due to the modifications made to the lipid A by the PhoPQ two component system. Host defense peptides (HDPs) are like colistin in that they are both positively charged and display amphipathic character, however are not impacted by lipid A modifications to the extent of colistin. To understand how HDPs can penetrate colistin resistant membranes we performed a deep mutational scanning analysis of protegrin-1 and revealed that amino acids mutations that resulted in alteration of peptide structure had more impact on antimicrobial activity than a reduction in charge. Probing single and double amino acid variants using membrane analysis and molecular modeling revealed the loss of antimicrobial activity correlated with decreased inner membrane leakage and pore modeling predicting decreased pore size,

## Introduction

Multidrug-resistant *Klebsiella pneumoniae* is responsible for a wide array of healthcare-associated infections throughout the globe [1–8]. More specifically, as of 2024, the World Health organization has declared carbapenem-resistant *K. pneumoniae* the top priority pathogen [9]. Colistin, a last resort antibiotic, has been used in place of carbapenems to treat carbapenem-resistant *K. pneumoniae* [2, 3, 7, 10–12]. The two *K. pneumoniae* clonal groups most associated with global carbapenem resistance are CG258 and CG15 [3, 13, 14]. While CG15 is more commonly associated with resistance to third-generation cephalosporins, as of 2017 CG258 was responsible for approximately 70% of carbapenem-resistant infections, with cases documented globally [3, 8]. Within this clonal group consists of three sequence types, ST258, ST512, and ST11. Of these sequence types, ST258 has been associated with the majority of carbapenem resistant outbreaks as well as global dissemination of carbapenemases [8, 13, 15, 16]. Importantly, we are now finding an increase in colistin resistance with ST258 clinical isolates [7, 17–20].

Polymyxin B and E (colistin) are cationic cyclic lipopeptides with a charge of +5 that bind to the lipopolysaccharides on the outer membrane of Gram-negative bacteria [10, 21–24]. More specifically, the positively charged diaminobutyric acid residues of colistin bind to the negatively charged phosphate groups on the lipid A portion of a lipopolysaccharide molecule (LPS), displacing the divalent cations already present. This allows the hydrophobic tail of colistin to penetrate the outer membrane, resulting in local membrane curvature. Eventually, colistin will form aggregates within the outer membrane, resulting in structural instabilities. These instabilities allow colistin to cross the lipid bilayer and access the inner membrane. At the inner membrane colistin will cause membrane thinning that will result in cytosolic leakage and bacterial cell death [10, 21–24]. Colistin resistance in *K. pneumoniae* has been largely attributed to the modifications on the outer membrane that alter the charge or structure of the lipid A portion its lipopolysaccharides (LPS) [10, 21–24]. The overall charge of the bacterial membrane can be increased by the replacement of negatively charged phosphates on the lipid A molecule by positively charged chemical groups such as phosphoethanolamine (PEtN) and 4-amino-4-deoxy-L-arabinose (L-Ara4N) [10, 21–24]. This will reduce colistin’s affinity for the bacterial cell due to the disruption of the initial electrostatic interaction. Furthermore, the acyl chains of a lipid a molecule can be altered to increase the rigidity of the bacterial membrane, resulting in the mechanistic expulsion of colistin and thereby resistance [22, 25]. The PhoPQ two-component system has been shown to be a global regulator implicated in colistin resistance in Gram-negative bacteria, including *K. pneumoniae* [26–29]

Host defense peptides are not equally affected by the modifications to the LPS that cause resistance to polymyxins [30–32]. Compared to colistin that directly targets lipid A to displace the stabilizing cations [10, 21–24], host defense peptides interact more generally with the LPS having multiple mechanisms of membrane disruption. These include the ‘detergent like’ carpet model or the formation of pores through the toroidal pore and barrel-stave models [33–36]. Previous work describing host defense peptide antimicrobial activity towards *K. pneumoniae* revealed that peptides with increased backbone steric hinderance had improved activity, in particular towards colistin-resistant ST-258 *K. pneumoniae* strain MKP103 [37, 38]. This suggests that not all host defense peptides are affected equally by the modifications providing colistin resistance and there are important physiochemical properties that allow them to retain antimicrobial activity towards colistin-resistant Gram-negative pathogens.

Here we describe the investigation into the strong antimicrobial activity of protegrin-1 towards colistin-resistant *K. pneumoniae* MKP103 through a multi-level analysis (**Figure 1**). Deep mutational peptide scanning revealed specific positions necessary for antimicrobial activity in protegrin-1. Furthermore, we found intricate changes to secondary structure could impact antimicrobial activity, while comparatively larger changes to the positive charge of the peptide were needed to impact activity. Specifically, within the triple arginine motif of protegrin-1 a proline mutation decreases antimicrobial activity at position 9 but not at position 11. Mutation of a second arginine residue within this region, regardless of the position of the proline resulted in complete loss of antimicrobial activity. Investigation into the peptide variants membrane interactions demonstrated changes in cytosolic leakage, which corresponded to the respective activity of our proline peptide variants. Moreover, despite a loss of activity we found an increase in outer membrane disruption with our inactive double mutation peptide variants. Molecular modeling revealed a distinct reduction in diameter and overall pore collapse of the modeled beta-sheet variant octameric pores as well as the rapid loss of beta-sheet character in the variants’ monomeric models. These findings challenge the currently set precedent of colistin resistance being largely attributed to differences in charge, where we see with protegrin-1 even with the cysteine bonds in place, intricate changes in structure can impact pore formation in colistin-resistant *K. pneumoniae*.

**Figure 1.**
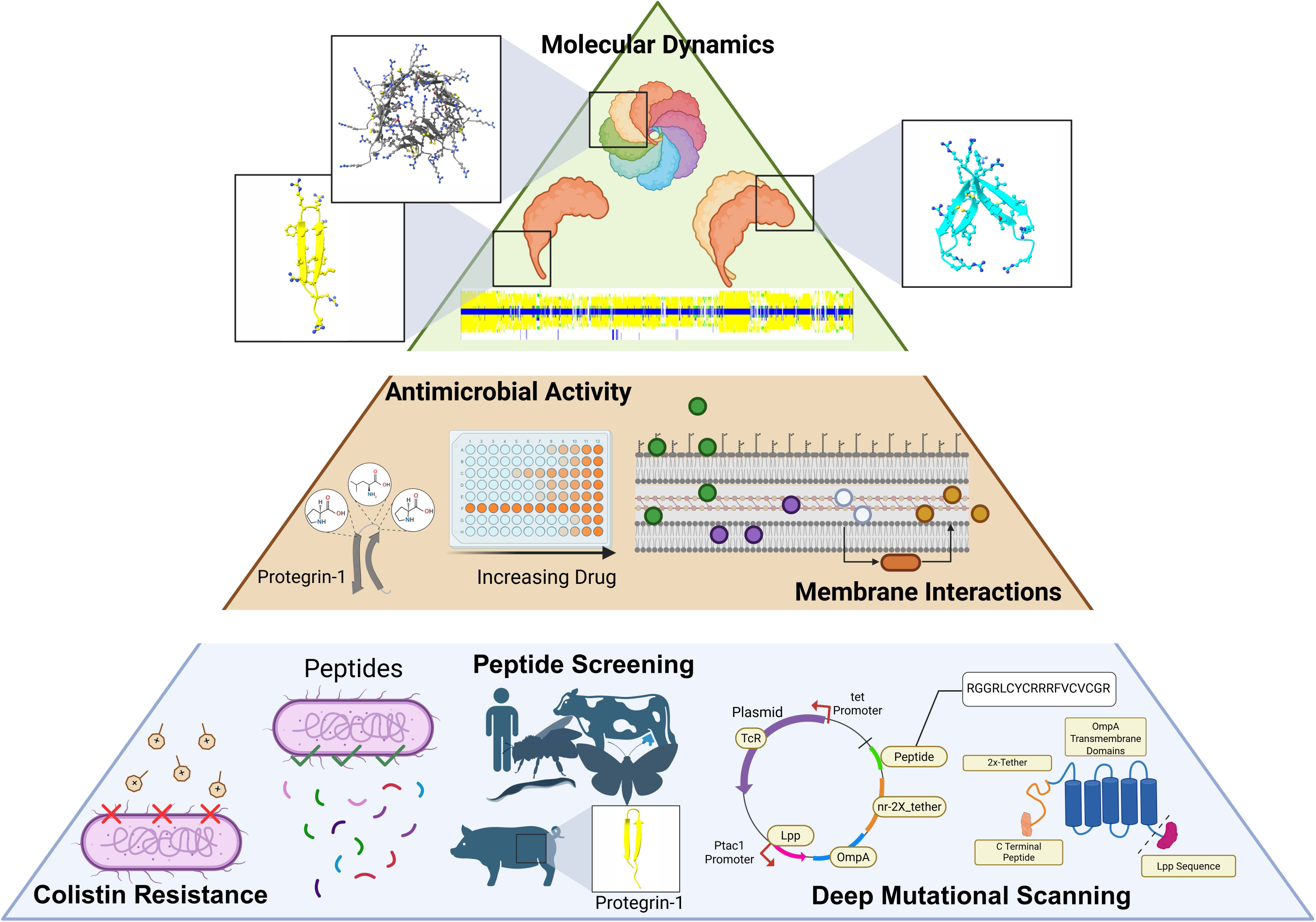
Multi-level analysis used to investigate how protegrin-1 can bypass colistin resistance. Our initial determination of MKP103 to have colistin-resistance yet has variable susceptibility to peptides of different structural class was used to inform our deep mutation scanning utilizing a genetic screening platform (bottom). We then investigate peptide variants mutated in different regions of the triple arginine motif for changes in antimicrobial activity, disruption to both the outer and inner membrane (middle). We ultimately use molecular modeling of peptides and octameric pores and combine this with molecular dynamic simulations to assess peptide structure stability (top). Octameric, dimeric and monomeric models were generated using Alpha Fold and Biorender was used for both vector production and for the creation of the final figure.

## Results

### Host defense peptides do not equally bypass *K. pneumoniae* MKP103 PhoP acquired polymyxin resistance

An increasing number of *K. pneumoniae* isolates are becoming resistant to the last resort antibiotic colistin [7, 17–20]. Colistin is positively charged and displays amphipathic structure, characteristics shared by host defense peptides [21, 39–41]. Host defense peptides are a stressor that is responded to by the PhoPQ two-component system, although the modifications to the LPS causing colistin resistance does not equally apply to all host defense peptides [30–32, 42]. To expand on our understanding of the varying impact of PhoPQ mediated colistin resistance on different host defense peptides first we validated *K. pneumoniae* MKP103 resistance to polymyxins was mediated by upregulation *phoP* and tested the impact of this resistance to antimicrobial activity of different classes of host defense peptides.

To validate PhoPQ mediated polymyxin resistance in *K. pneumoniae* MKP103, we first identified the minimal inhibitory concentration (MIC) of polymyxin B and colistin (polymyxin E) next to other well published *K. pneumoniae* strains. We found minimal polymyxin resistance with hypervirulent *K. pneumoniae* NTUH [43] and KPPR1S [44], drug-resistant *K. pneumoniae* ATCC 700603 [45], and type strain *K. pneumoniae* ATCC 13883 [46] (MICs < 1 µmol L^-1^) when compared to *K. pneumoniae* MKP103, a genetically modified ST-258 strain used for the creation of a transposon library (MIC 27 µmol L^-1^) [47, 48] (**Table S1, Figure 2A**). We then determined the baseline expression levels of response regulator *phoP* with RT-qPCR to compare to their respective polymyxin resistance [49, 50] and found high levels of *phoP* expression correlated with colistin resistance in *K. pneumoniae* MKP103 (**Figure 2A,Figure S1**). Therefore, we determined *K. pneumoniae* MKP103 to be the ideal screening candidate for our investigation into PhoPQ mediated colistin resistance.

**Figure 2.**
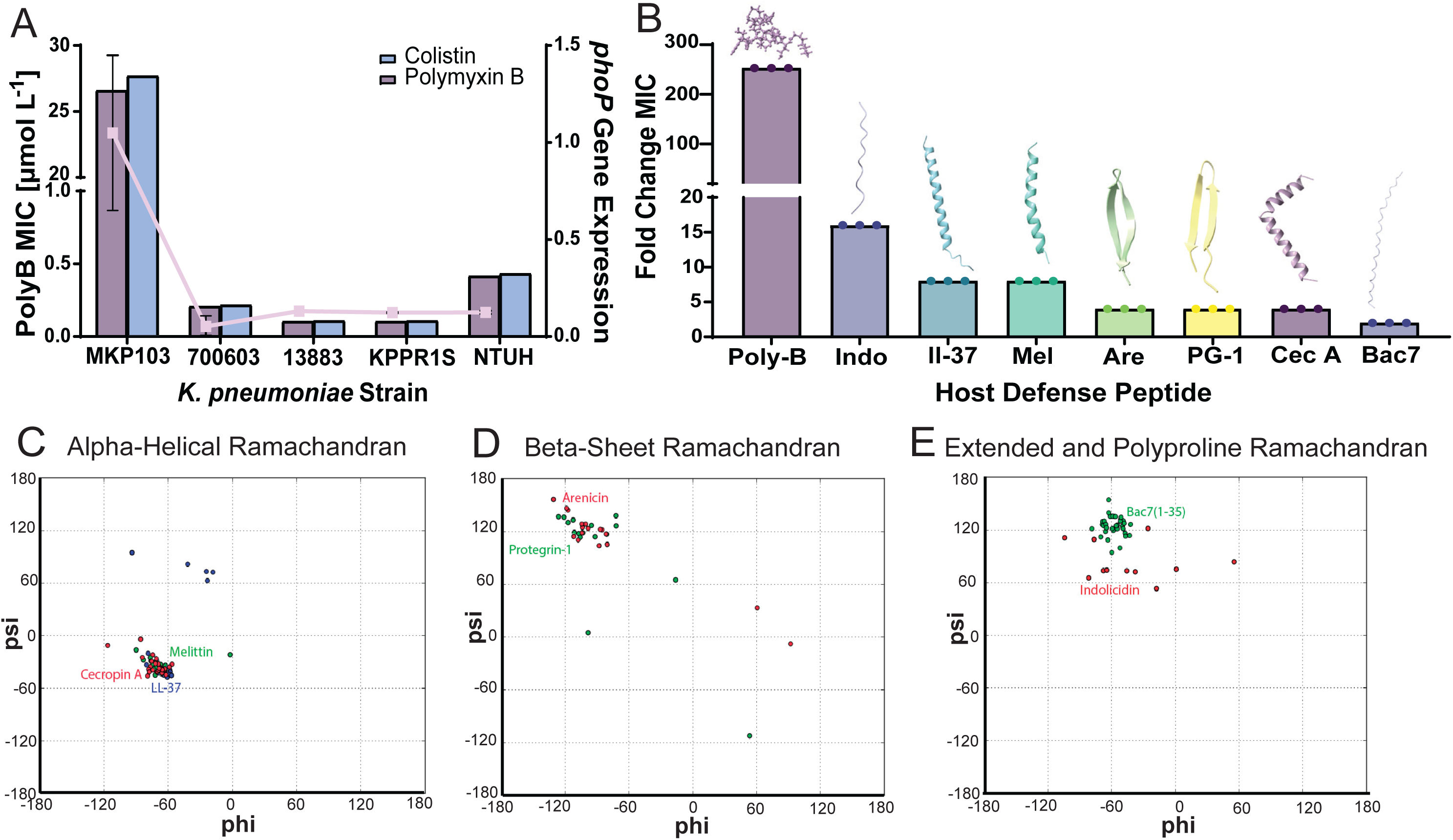
Host defense peptides do not equally bypass *K. pneumoniae* MKP103 PhoP acquired polymyxin resistance. This figure shows bacteria and peptide screening to determine ideal candidates to investigate colistin resistance. Colistin and polymyxin B MICs against drug-resistant *K. pneumoniae* MKP103, hypervirulent *K. pneumoniae* NTUH and KPPR1S, drug-resistant *K. pneumoniae* ATCC 700603, and type strain *K. pneumoniae* ATCC 13883 (bars) overlayed with the baseline *phoP* expression in exponential growth phase, normalized to housekeeping gene *rpoD* and graphed in triplicate with error shown as ±SEM (Figure 2A). Fold change in MICs between the *K. pneumoniae* MKP103 parental isolate and a *ΔphoP* transposon mutant (KPNIH1_10030-701::T30) of polymyxin B (Poly-B), indolicidin (Indo), Melittin (Mel), Arenicin (Are), protegrin-1 (PG-1), and CecropinA (Cec A) (Figure 2B). MIC assays were performed in triplicate for the data shown. Figures 2C**-2E** show the Ramachandran plots depicting the psi and phi angles of alpha helical peptides, beta sheet peptides, and extended/polyproline peptides, respectively.

To determine a candidate host defense peptide for our deep mutational scanning, we used the parental MKP103 and its *ΔphoP* transposon mutant (KPNIH1_10030-701::T30) and compared MICs of different structural classes of antimicrobial peptides (**Table 1**). As shown with previous studies on colistin resistance [30–32], we found that PhoP mediated polymyxin resistance does not extend equally to host defense peptides compared to polymyxin B and E, which had a 256-fold change between the parental strain and *ΔphoP* mutant (**Figure S1, Figure 2B**). Specifically, the largest fold change in MIC between the parental strain and *ΔphoP* mutant was with indolicidin (16-fold), an extended host defense peptide with a random coil structure (**Figure 2B**). Furthermore, when testing alpha-helical peptides, we saw LL-37 and melittin (8-fold change MIC) were similarly affected by PhoP mediated resistance while cecropin A was slightly less affected (4-fold change MIC). When testing beta-sheet peptides, protegrin-1 and arenicin both had a 4-fold change in MIC between the parental strain and *ΔphoP* mutant. Finally, as expected due to its internal mechanism of killing [51], the polyproline peptide bac7 (1-35) had only 2-fold change in MIC between the MKP103 parental strain and *ΔphoP* mutant. For a comparison of the peptide backbone steric hindrance of the tested host defense peptides following up on previously observed finding [37], we created Ramachandran plots of the host defense peptides displaying the dihedral rotation angles of each amino acid residue. The alpha helical host defense peptides (ll-37, cecropin, and melittin) plotted within the right-handed alpha-helical region of the Ramachandran plot (**Figure 2C**) revealing the precise angles needed to form a helical protein [52–54] due to the flexibility of the peptide backbone. The beta-sheet peptides (protegrin-1 and arenicin) and extended peptides (bac7 (1-35) and indolicidin) plotted within the beta-sheet/collagen triple helix region of the Ramachandran plot (**Figure 2D, and 2E**), a region known for being the most restrictive for amide bond rotation [54]. Although indolicidin localized within this region, the individual amino acids displayed a range of flexibility, shown by the spread on the Ramachandran plot. Collectively, host defense peptides that had tight localization to the beta-sheet/collagen triple helix region of the Ramachandran plot were less impacted by *K. pneumoniae* MKP103 PhoP mediated colistin resistance. Protegrin-1, along with having a low fold change between parental MKP103 and *ΔphoP* mutant, also has a well characterized structure and mechanism of action [55–57]. Furthermore, this peptide’s two disulfide bonds, which are responsible for protegrin-1’s tightly constraint structure, are ideal for molecular dynamic simulations [58]. For these reasons we chose this peptide as our candidate for deep mutational scanning to investigate the ability of this host-defense peptides to bypass *K. pneumoniae* colistin resistance.

**Table 1.**
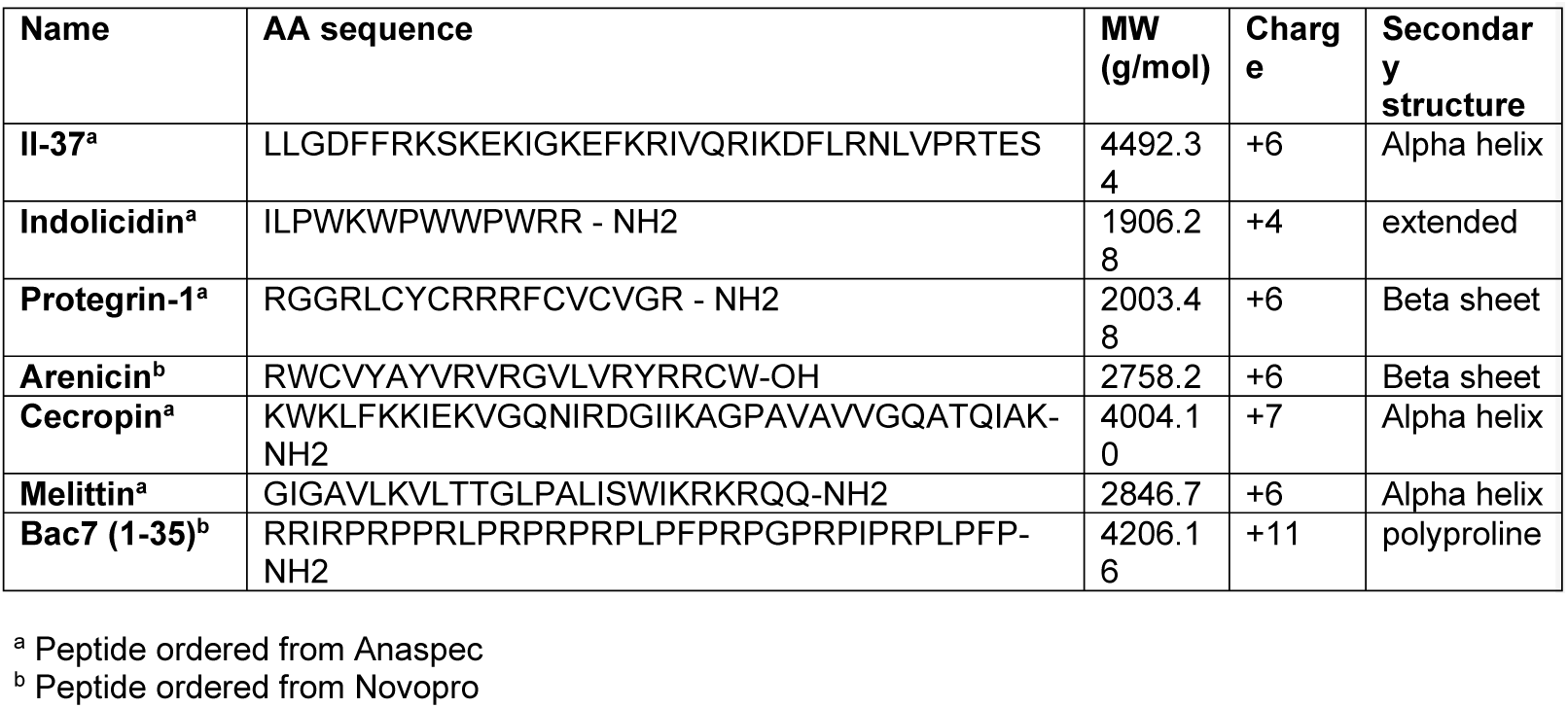
Host defense peptides tested for antimicrobial activity towards *K. pneumoniae* MKP103 and MKP103 *ΔphoP* mutant.

### Protegrin-1 deep mutational scanning library shows reveals charge and structure are important for antimicrobial activity towards colistin-resistant *K. pneumoniae* MKP103

To understand the importance of individual amino acids to the antimicrobial activity of beta-sheet peptide towards colistin-resistant *K. pneumoniae* we next aimed to perform deep mutational scanning analysis of the beta-sheet peptide protegrin-1. For this we utilized the Surface Localized Antimicrobial Display (SLAY) system that was designed to display antimicrobial peptides on the surface of Gram-negative bacteria to capitalize on the high-throughput advantages of next-generation sequencing [59]. Further optimization of this platform led to the development of deep mutational scanning with SLAY (dmSLAY), using this tool to assess protegrin-1 and describe features driving membrane selectivity between bacteria and mammalian cells [60]. As previously described, the creation of the protegrin-1 dmSLAY ensured a high level of conservation was given to the mixed base primer sequence (**Table S2, Figure S2**), with the possibility of 7-12 changes at each amino acid residue, resulting in 1-9 mutations per peptide variant. We aimed to expand on this impactful study by testing the protegrin-1 dmSLAY library in colistin-resistant *K. pneumoniae* MKP103.

We collected the plasmid-based library from colistin-resistant *K. pneumoniae* MKP103 following growth for four hours with and without plasmid induction with isopropyl β-D-thiogalactoside (IPTG). To predict antimicrobial activity of the peptide variants in our library, we used next generation sequencing to quantify the plasmids from the induced versus the uninduced population **(Table S3)**. Sequences that express for an active peptide variant are removed from the population of plasmids following induction resulting in a negative log2 fold change in reads. With the goal to understand what amino acids were implicated in retaining the antimicrobial activity of protegrin-1, we looked for retention within the population of peptide variants rather than removal and used a log2 fold change value of 0 or higher is indicative of lack of antimicrobial activity. Overall, we identified approximately 1700 peptide sequence variants lacking activity against *K. pneumoniae* MKP103 **(Figure 3A; Supplemental data file 1).** To determine which specific amino acid residues were necessary to bypass colistin resistance, we chose to investigate individual mutations that occurred most often in our inactive peptide variant population, rather than testing peptide variants from the screen **(Table S4)**. We had some individual amino acid mutations that occurred only ≤20 peptide variants. However, we were intrigued to find several amino acid mutations that were in the majority of peptide variants retained in the population **(Table 2**, **Figure 3B).** We chose not to investigate mutations with less than 25 occurrences, those that did not involve an amino acid class change, and those that impacted disulfide bonds, as literature has already shown their importance for protegrin-1’s antimicrobial activity [61, 62]. This resulted in the selection of five individual mutations in different structural regions of protegrin-1 to investigate their effects alone on the surface display system (**Figure 3B**). Specifically, the glycine at position to mutated to an arginine (G2R) increasing the charge of the unstructured tail region was found 316 times within the population of peptide variants that had a log2 fold change >0. A leucine mutation to an isoleucine (L5I) and phenylalanine to leucine (F12L) within the beta-sheet region were found in 218 and 834 of peptide variants retained in the population, respectively. Finally, we found mutations in 2 arginine residues of the triple arginine loop region (R9P and R10L) times 295 and 299 within the population of peptide variants retained in the population, respectively and these variants most often occurred together.

**Figure 3.**
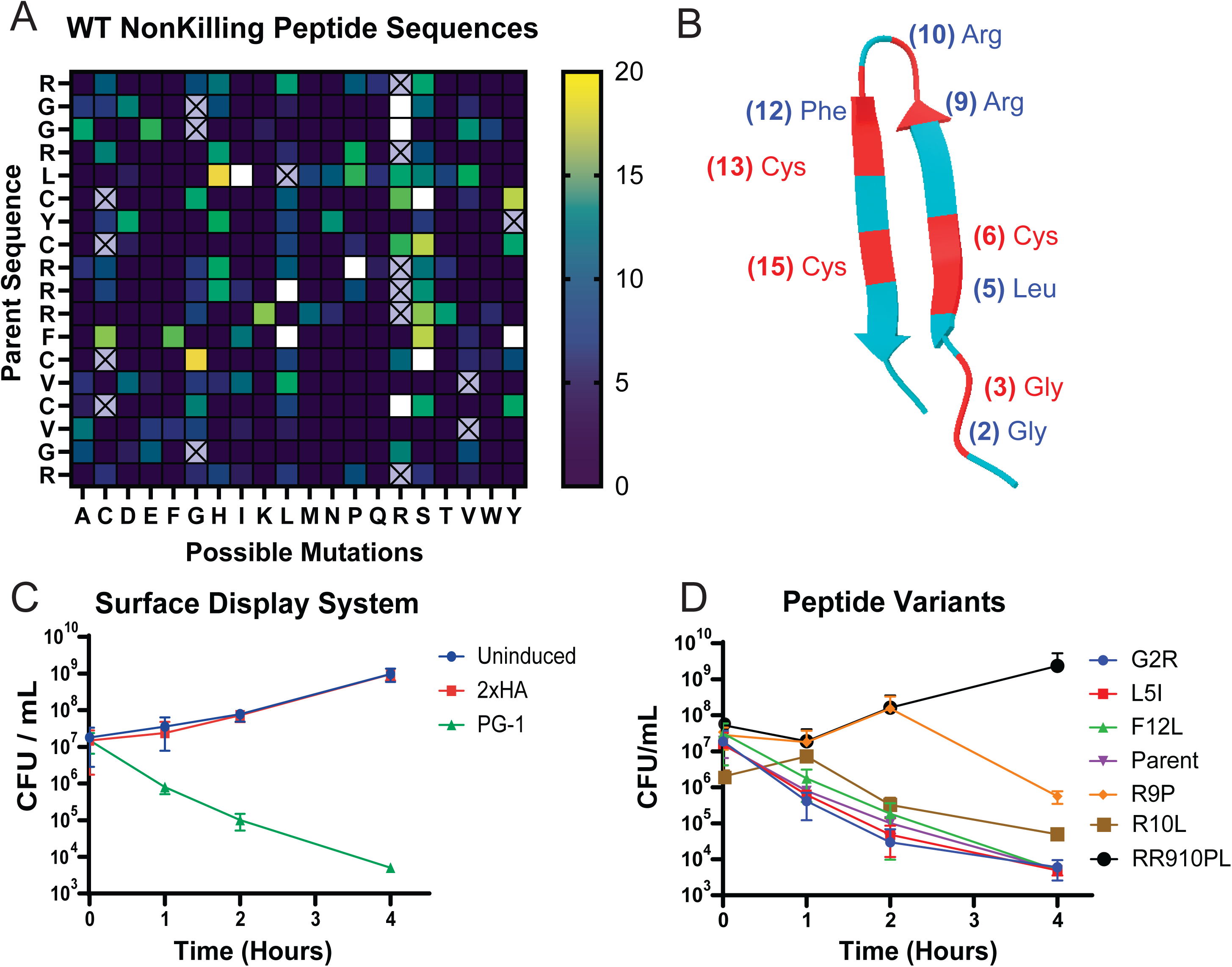
Protegrin-1 deep mutational scanning library shows reveals charge and structure are important for antimicrobial activity towards colistin-resistant *K. pneumoniae* MKP103. This figure shows the results of the initial SLAY screening as well as validation with direct cloning. Figure 3A is a heat Map of SLAY library analysis showing the number of occurrences of the amino acid mutations in our Illumina sequencing populations retained with induction with the parent peptide sequence on the y axis and possible mutations on the x axis. White boxes denote mutations outside of the defined maximum range of the heat map (20 occurrences). Figure 3B shows the Alpha Fold representation of protegrin-1 peptide with the mutations identified from the screen as outside of the defined maxima show with their location highlighted in red and amino acid position number in parentheses. The amino acid mutations denoted in blue text were cloned in the surface display system for validation by enumeration of CFU mL^-1^ following IPTG plasmid induction over four hours with display of eighth the parent peptide protegrin-1, uninduced plasmid control, and negative control 2xHA (Figure 3C) and peptide variants (Figure 3D) identified in the screen. The surface display validation assays were performed in triplicate with error shown.as as ±SEM.

**Table 2.** Occurrence of amino acid mutations in the peptide variant sequences lacking antimicrobial activity.

| Amino Acid Mutation | Amino Acid Class Change | Number of Occurrences |
| --- | --- | --- |
| F12L | non-polar to non-polar | 834 |
| G2R | non-polar to positive-charge | 316 |
| R10L* | positive-charge to non-polar | 299 |
| R9P* | positive-charge to non-polar | 295 |
| L5I | non-polar to non-polar | 218 |
| C6S | non-polar to polar | 35 |
| C13S | non-polar to polar | 27 |
| C15R | non-polar to positive-charge | 22 |
| G3R | non-polar to positive-charge | 21 |
| F12Y | non-polar to polar | 21 |

\*Amino acid mutations found together in 89% occurrences

To test the selected individual mutations using the SLAY platform, we cloned the single amino acid variants of protegrin-1 into the genetic screening system **(Table S5, S6)** and induced the plasmid in colistin-resistant *K. pneumoniae* MKP103 over a four-hour period enumerating CFU mL^-1^. When testing the parental peptide displayed on the surface with 1 mM IPTG induction we found there was a 5-log reduction in CFU mL^-1^ when compared to an uninduced plasmid control, and bacteria harboring the system with a tandem influenza hemagglutinin peptide (2xHA) as a negative control **(Figure 3C)**. Testing the individual amino acid mutations next to the parental peptide, we found the mutation of an arginine to leucine (R10L), glycine to arginine (G2R), leucine to isoleucine (L5I), and phenylalanine to leucine (F12L) mutations displayed minimal change in CFU mL^-1^ compared to the surface display of the parental peptide (**Figure 3D).** However, when introducing a proline mutation at position 9 (R9P), we saw this single mutation could reduce the activity of the peptide towards colistin-resistant *K. pneumoniae* MKP103 by 2-fold compared to our parental peptide. Intriguingly, we found the arginine to proline mutation in our next-generation sequencing occurred approximately 89% of the time with an arginine to leucine mutation at position 10 (R9P, R10L). Therefore, we also tested these amino acid mutations together using SLAY and found the complete loss of activity of the peptide with two arginine amino acid residues mutated within the triple arginine motif (**Figure 3D**). Overall, our deep mutational scanning of protegrin-1 identified candidate amino acids that are important for antimicrobial activity towards colistin-resistant *K. pneumoniae.* Although many of these mutations needed additional mutations for a reduction in antimicrobial potential, we identified a single arginine to proline mutation at position 9 within the beta-sheet to turn transition region of the triple arginine motif that alone could impact antimicrobial activity and the complete loss of activity when this was accompanied by a second mutation within this region.

### The arginine to proline mutation impedes antimicrobial activity only in the beta-sheet to turn transition region of the protegrin-1 arginine loop

With the intriguing findings from our deep mutational scanning of protegrin-1 arginine loop beta-sheet to turn transition region, we had the single (R9P and R10L) and double (R9P, R10L) protegrin-1 variants synthesized to test their antimicrobial activity towards of *K. pneumoniae* MKP103 parental strain next to the *ΔphoP* mutant, mutants downstream of PhoP regulon, and a panel *K. pneumoniae* clinical isolates. In addition to testing amino acids discovered in our dmSLAY analysis, we also synthesized a single and double variant with a proline mutation at position 11 (R11P and R10L, R11P). This placed the proline amino acid substitution within the peptide’s turn region rather than its beta-sheet to turn transition region **(Table 3)** and test the importance of the changes to peptide positive charge versus the secondary structure that were introduced by the R9P mutation. Furthermore, in order to investigate the position itself rather than the amino acid mutation found in that position, we also synthesized alanine and lysine single peptide variants at positions 9 (R9A and R9K) and 11 (R11A and R11K).

**Table 3.**
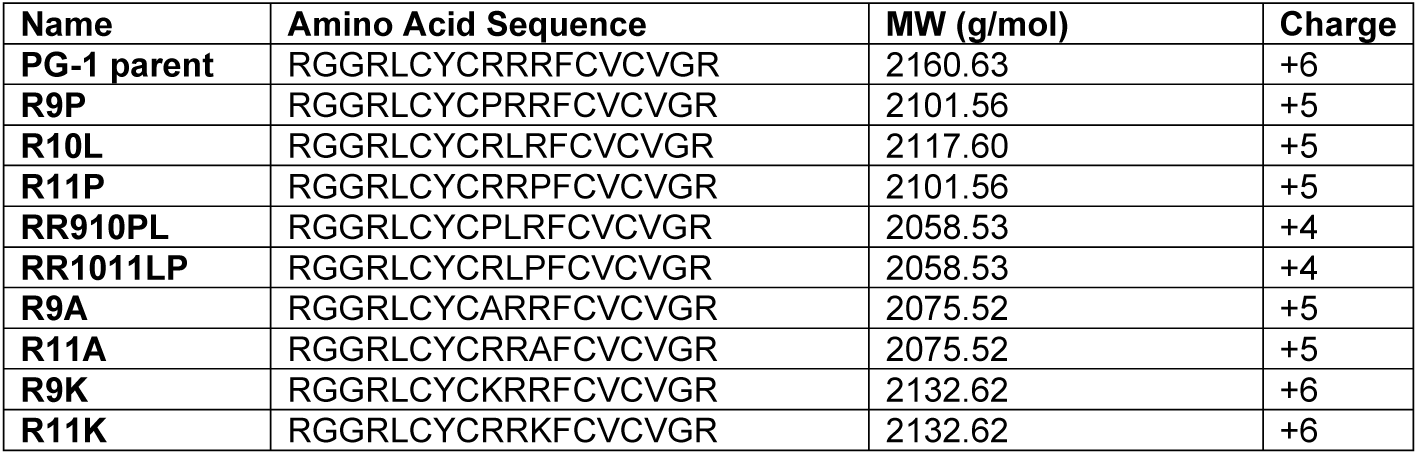
Synthesized Peptide Variants.

When testing antimicrobial activity towards the colistin-resistant *K. pneumoniae* MKP103 used for screening, both the R9P and R10L variant displayed a reduction of activity compared to protegrin-1 (MIC 15.23 and 15.11 µmol L^-1^, respectively) **(Table 4)**. We observed that despite a reduction in charge (charge +5), the R11P variant retained the same level of antimicrobial activity toward our colistin-resistant isolate as protegrin-1 (charge +6) (MICs 3.8 µmol L^-1^ respectively). When testing the double mutant variants (R9P, R10L and R10L, R11P) (charge +4), we found they both lost activity towards *K. pneumoniae* MKP103 (MIC >30 µmol L^-1^). Interestingly, when substituting either position 9 or 11 with an alanine (R9A or R11A) we found the similar antimicrobial activity to the R9P peptide variant (MICs 15.41,15.41, 15.23 µmol L^-1^, respectively). However, when substituting position 9 and 11 with a lysine (R9K or R11K) we have smaller changes in activity, indicating charge retention at these positions has variable impacts on antimicrobial activity (MICs 3.75 and 7.50 µmol L^-1^, respectively).

**Table 4.** Minimum inhibitory concentration of protegrin-1 and peptide variants towards *K. pneumoniae* MKP103 and transposon mutants.

| Peptide Variant | MIC ($\mu\text{mol L}^{-1}$ / $\mu\text{g mL}^{-1}$) | | | | | |
| --- | --- | --- | --- | --- | --- | --- |
| | MKP103 WT | $\Delta phoP$ | $\Delta pagP$ | $\Delta pagL$ | $\Delta arnT$ | $\Delta pmrB$ |
| **PG-1 parent** | 3.70/8 | 0.12/0.25 | 3.70/8 | 3.70/8 | 3.70/8 | 3.70/8 |
| **R9P** | 15.23/32 | 0.48/1 | 15.23/32 | 15.23/32 | 15.23/32 | 15.23/32 |
| **R10L** | 15.11/32 | 1.89/4 | 15.11/32 | 15.11/32 | 7.56/16 | 7.56/16 |
| **R11P** | 3.81/8 | 0.95/2 | 0.95/2 | 0.95/2 | 1.90/4 | 1.90/4 |
| **RR1011LP** | 31.09/64 | 0.97/2 | 15.55/32 | 15.55/32 | 15.55/32 | 15.55/32 |
| **RR910PL** | 31.09/64 | 3.89/8 | 31.09/64 | 31.09/64 | 31.09/64 | 31.09/64 |
| **R9A** | 15.41/32 | 0.96/2 | 1.93/4 | 3.85/8 | 1.93/4 | 3.85/8 |
| **R11A** | 15.41/32 | 0.48/1 | 0.96/2 | 0.96/2 | 0.96/2 | 1.93/4 |
| **R9K** | 3.75/8 | 0.47/1 | 0.94/2 | 0.94/2 | 0.94/2 | 0.94/2 |
| **R11K** | 7.50/16 | 0.47/1 | 0.94/2 | 0.94/2 | 0.94/2 | 0.94/2 |

To broaden the impact of our findings beyond *K. pneumoniae* MKP103 we selected colistin-resistant clinical isolates from the Mult-drug-Resistant Organism Repository and Surveillance Network (MRSN) *K. pneumoniae* diversity panel to test the peptide variants against [63]. We found although the majority of isolates (64 MRSN isolates) were colistin sensitive (MIC ≤ 0.87 µmol L^-1^), there were 31 isolates that displayed moderate colistin resistance (MIC 1.73-6.92 µmol L^-1^), and 5 isolates (MRSN 13768, 430414, 430405, 560539, and 728987) that displayed elevated colistin resistance (MICs >13 µmol L^-1^) **(Table S7)**. Of note, two isolates, MRSN 13768 and 430414, had colistin MICs comparable to that of *K. pneumoniae* MKP103 (27.69 µmol L^-1^). However, when assessing our protegrin-1 peptide variants towards these clinical isolates, we found that majority our MICs were relatively similar to the parental peptide protegrin-1, with the R9P single variant displaying the least antimicrobial activity (3.81 µmol L^-1^) (**Figure 4A**). When testing the double mutant variants (R9P, R10L and R10L, R11P) we found they lost antimicrobial activity towards 3 of the 5 colistin resistant MRSN clinical isolates. Specifically, MRSN 430405 (colistin MIC 27.69 µmol L^-1^), was completely resistant to the beta turn double mutant peptide variant (R10P, R11L MIC 31.09 µmol L^-1^) but not the beta sheet double variant (R9P, R10L MIC 7.77 µmol L^-1^). Furthermore, MRSN 560539 (colistin MIC 13.85 µmol L^-1^) displayed the same level of resistance to both beta sheet and beta turn double variants (MICs 15.54 and 15.55 µmol L^-1^, respectively) and MRSN 728987 (colistin MIC 13.85 µmol L^-1^) was slightly more sensitive to the double mutant peptide variants (MIC 7.77 µmol L^-1^).

**Figure 4.**
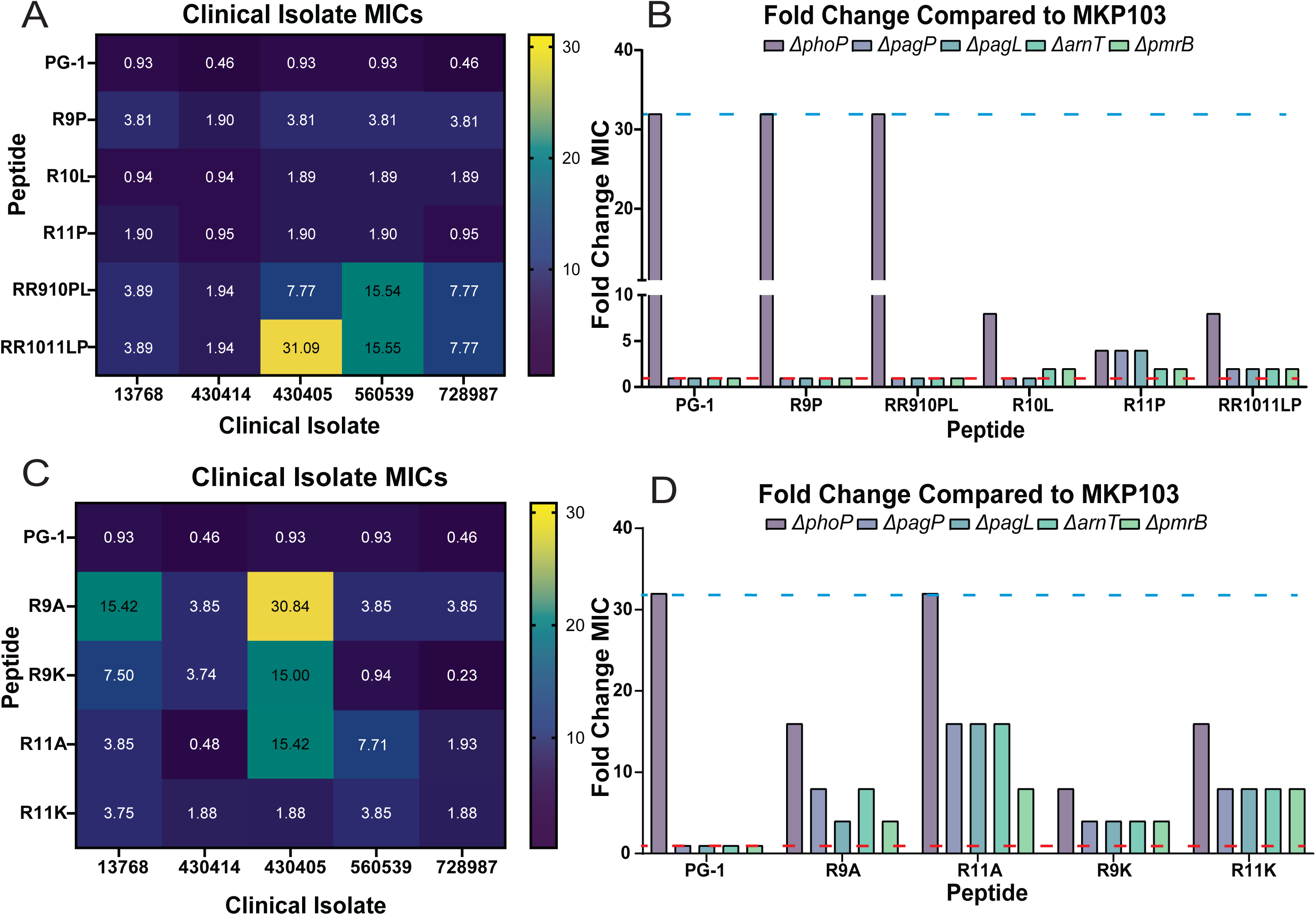
The arginine to proline mutation impedes antimicrobial activity only in the beta-sheet to turn transition region of the protegrin-1 arginine loop. The figures show MICs (µmol L^-1^) of synthesized peptide variants and fold-change in MICs between colistin-resistant *K. pneumoniae* MKP103 and transposon mutants lacking important LPS modifying genes. Figure 4A shows synthesized peptide variants were tested against five colistin resistant clinical isolates. Figure 4B shows the fold change of peptide MICs between the parental MKP103 and the *ΔphoP, ΔpagP, ΔpagL, ΔarnT,* and *ΔpmrB* transposon mutants (mut/WT). Figure 4C and 4D shows the alanine and lysine peptide variants MICs against the five colistin resistant clinical isolates and the fold change MICs between the parental MKP103 and transposon mutants. Lines are drawn on figures 4B and 4C follow the protegrin-1 parental peptide fold-change in MIC for *ΔphoP* (blue) and downstream genetic mutants (red). Peptides were all tested at a maximum concentration of 31 µmol L^-1^. MIC assays were performed in triplicate for the data shown.

When assessing the changes in MIC in the absence of PhoP regulon induction using the *ΔphoP* mutant, we found the beta sheet peptide variants (R9P and R9P, R10L) had a larger fold-change in antimicrobial activity between the mutant and parental *K. pneumoniae* MKP103 than the beta turn variants (R11P and R10L, R11P) **(Figure 4B)**. Of note, when comparing this the parental peptide protegrin-1, we found the beta sheet peptide variants (R9P and R9P,R10L) displayed the same 32-fold decrease in MIC toward the *ΔphoP* mutant as protegrin-1, whereas the beta turn peptide variants (R10L, R11P, and R10L, R11P) only displayed an 8-, 4-, and 8-fold decrease in MIC towards the *ΔphoP* mutant, respectively. Furthermore, we tested *K. pneumoniae* MKP103 transposon mutants of downstream genes within the PhoP regulon, involved in modifying either the rigidity or charge of the LPS [25, 42]. Specifically, PagP transfers a palmitate group to a lipid A molecule and PagL removes an acyl group, altering the membrane’s fluidity and permeability [25, 64], while ArnT and PmrB are involved in the addition of a 4-amino-4-deoxy-L-arabinose (L-Ara4N) on lipid A, increasing the positive charge of the outer membrane [25]. We found no difference between the MICs of parental protegrin-1, the beta sheet peptide variants (R9P and R9P, R10L), indicating a combination of LPS modifications are necessary for the sensitivity displayed by the *ΔphoP* mutant (**Figure 4B**). Interestingly, although the beta turn peptide variants did not show as much of a fold-change in MICs with the *ΔphoP* mutant as the beta sheet peptide variants and parental peptide protegrin-1, we found a slight fold-change increase in MIC between the downstream PhoP regulon mutants compared to the parental *K. pneumoniae* MKP103. Specifically, when testing the R11P variant, the *ΔpagL and ΔpagP* transposon mutants (KPNIH1_07455-208::T30 and KPNIH1_13050-101::T30, respectively) displayed 4-fold decrease in MIC (MIC 0.96 µmol L^-1^) **(Table S5)** compared to *K. pneumoniae* MKP103 (MIC 3.81 µmol L^-1^), while the *ΔarnT and ΔpmrB* transposon mutants (KPNIH1_24880-114::T30 and KPNIH1_08200-303::T30) displayed 2-fold decrease in MIC (MICs 1.90 µmol L^-1^). Finally, the beta turn double mutant peptide variant (R10L, R11P) all four PhoP regulon transposon mutants displayed a 2-fold increase in MIC.

Upon investigation of positions 9 (beta sheet) and 11 (beta turn) within protegrin-1 with alanine and lysine amino acid substitutions, we find much more variability within our colistin resistant clinical isolates than the mutations that came from the dmSLAY analysis of *K. pneumoniae* MKP103 **(Figure 4C)**. Specifically, alanine substitution at position 9 (beta turn) had MICs of 15.42 and 30.84 µmol L^-1^ towards MRSN 13768 (colistin MIC 27.79 µmol L^-1^) and MRSN 430505 (colistin MIC 13.85 µmol L^-1^), respectively. MRSN 430505 also displayed higher resistance towards the peptide variant with alanine substituted in the turn region (R11A MIC 15.00 µmol L^-1^) and lysine substituted in the sheet region (R9K MIC 15.42 µmol L^-1^). These peptides also displayed less variability between the antimicrobial activity towards the parental strain *K. pneumoniae* MKP103 and the PhoPQ regulon transposon mutants. Contrary to our variants identified from the initial screening, we found increased fold change in activity between the parental strain and PhoP regulon transposon mutants **(Figure 4D)**. Interestingly, we see these less bulky substitutions at position 11 variants (R11A, R11K) were more impacted by the downstream individual PhoP regulated changes than the position 9 variants (R9A, R9K). Specifically, R11A peptide variant when tested against the *ΔpagP, ΔpagL, and ΔarnT* transposon mutants resulted in a 16-fold decrease in MIC compared the parental strain, and *ΔpmrB* transposon mutant had an 8-fold lower MIC. When testing our peptide variants against other Gram-Negative bacteria we find that regardless of variant, *Burkholderia thailandensis* [65] exhibits complete resistance, while *Acinetobacter baumannii* [66] and both *Escherichia coli* [60, 67, 68] strains display high sensitivity to all peptide variants, suggesting these peptide variants identified in this study are unable to bypass Gram-negative bacteria with high levels of colistin resistance **(Table S8)**. Collectively, we found when proline was introduced to the turn region (R11P) of the triple arginine motif had less impact on the antimicrobial activity than when placed in the beta-sheet to turn transition region (R9P) and that removing an additional arginine from the triple arginine motif regardless of the proline position completely abolished antimicrobial activity towards colistin-resistant *K. pneumoniae* MKP103.

### Protegrin-1 double mutant variants with reduced charge display greater disruption of colistin-resistant outer membrane

We found modifications to the outer membrane LPS by the PhoP regulon increase resistance to our beta sheet peptide variants more than our beta turn variants, indicating a role in that amino acid in bypassing colistin resistance LPS modifications. Furthermore, we found a dichotomy between colistin resistance and peptide variant resistance when testing the MRSN clinical isolates, where the moderately elevated colistin resistant clinical isolates (MIC 1.73-6.92 µmol L^-1^), are more resistant to the double mutant peptide variants compared to the more colistin resistant clinical isolates (MIC 13 µmol L^-1^). Therefore, when assessing the mechanism of action of the peptide variants compared to the protegrin-1 parental peptide, we wanted to first probe the outer membrane integrity following treatment as protegrin-1 must interact and bypass the outer membrane before forming a pore at the inner membrane. To evaluate changes in outer membrane permeability, we used N-Phenyl-1-naphthylamine (NPN), a hydrophobic small molecule impermeable to an intact outer membrane that fluoresces weakly with increased fluorescence observed upon outer membrane disruption [69–71].

When testing the parental strain *K. pneumoniae* MKP103, our single point mutations (R9P, R10L and R11P) at 31 µmol L^-1^ behave similarly to protegrin-1 but also polymyxin B (MIC 27 µmol L^-1^) **(Figure 5A).** Unexpectedly, the double mutation variants (R9P, R10L and R10L, R11P) when tested at 31 µmol L^-1^ display higher fluorescence of NPN than parent protegrin-1, indicating increased disruption of the outer membrane of colistin-resistant *K. pneumoniae* MKP103 **(Figure 5B).** Interestingly, when testing our *ΔphoP* mutant **(Figure 5C and 5D)** and downstream mutants modifying both charge and fluidity, all peptide variants display higher fluorescence of NPN (**Figure S3)**. Our beta sheet double peptide variant in these cases had activity similar to or less than our parent peptide. Comparing the endpoint fluorescence across isolates we find the least amount of disruption from our single mutant peptide variants with our *ΔpagL* transposon mutant **(Figure 5E)**. We also find that in every isolate our more active variant (R11P) is less effective at disrupting the outer membrane compared to our less active variant (R9P). When observing endpoint fluorescence of our double mutant variants we also see the least amount of disruption with our *ΔpagL* transposon mutant **(Figure 5F)**. When testing our most colistin resistant clinical isolates, the peptide variants also display higher fluorescence of NPN **(Figure S4)**. However, our beta turn double peptide variant (R10L, R11P) disrupts the outer membrane more effectively than the beta sheet double peptide variant (R9P, R10L) **(Figure S4)**. Overall, we account for some but not all of the changes in antimicrobial activity towards parental MKP103 with differences in peptide interactions within the outer membrane.

**Figure 5.**
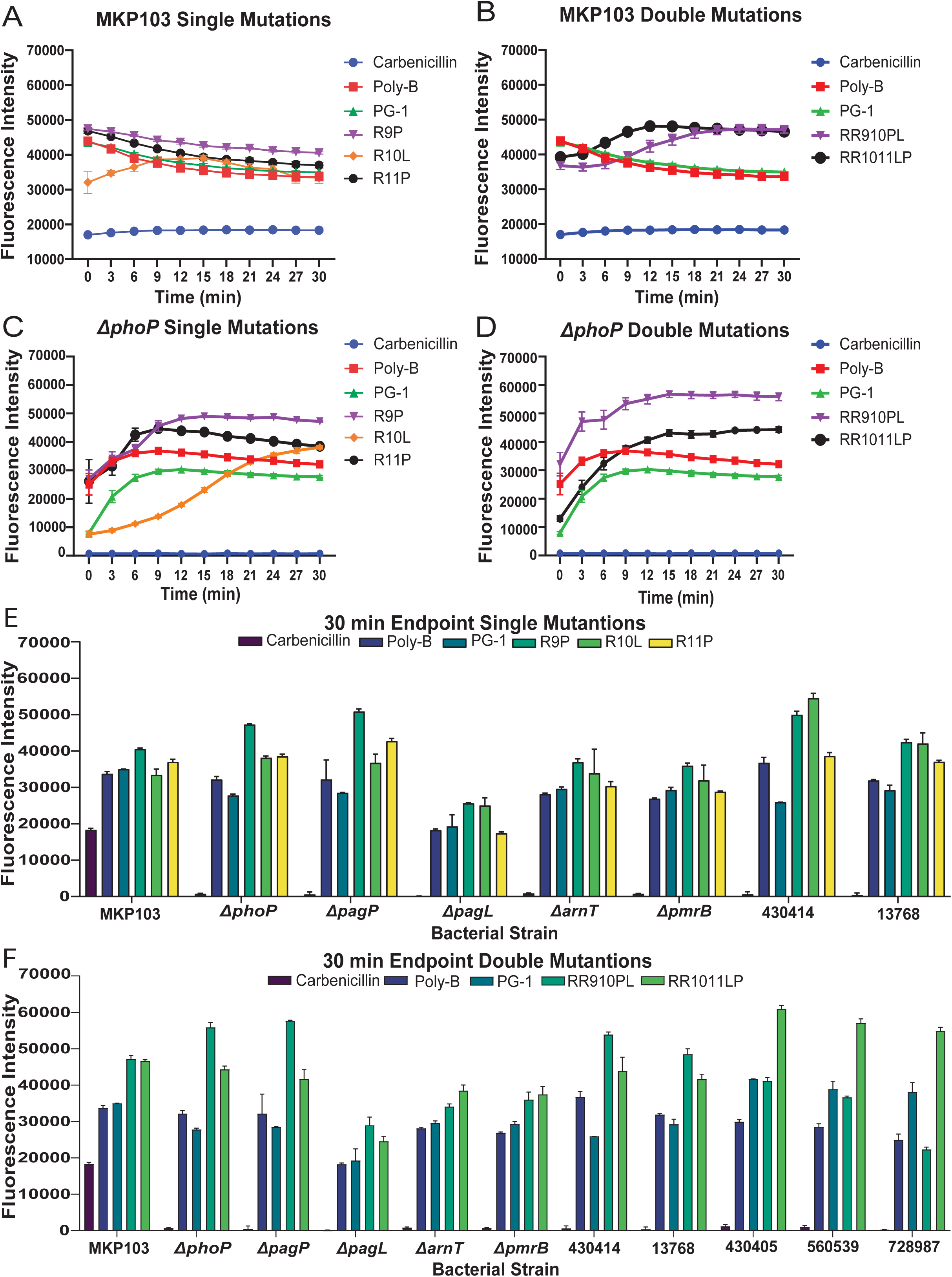
Protegrin-1 double mutant variants with reduced charge display greater disruption of colistin-resistant outer membrane. Outer membrane fluorescence was monitored over 30 minutes with peptide variants and controls carbenicillin and polymyxin B. Figure 5A shows *K. pneumoniae* MKP103 normalized NPN fluorescence after treatment with single peptide variants and Figure 5B shows the double peptide variants. The NPN fluorescence for the *ΔphoP* transposon mutant is shown for peptide variants in Figures 5C **and 5D**. Endpoint fluorescence reads with single peptide variant mutants with top five colistin resistant MRSN isolates, parental MKP103, *ΔphoP, ΔpagP, ΔpagL, ΔarnT, and ΔpmrB* transposon mutants is shown on Figure 5E, and double peptide variant mutants on Figure 5F. All peptides and controls were tested in triplicate at a concentration of 31 µmol L^-1^ with error shown as ±SEM.

### The arginine to proline mutation in the beta-sheet transition limits cytosolic leakage in single and double mutation variants

Protegrin-1 has a membrane disruptive mechanism of action and has been described to form pores in the inner membrane of the bacteria [55, 72]. To assess the peptide variants loss of membrane disruptive killing, we investigated their ability to depolarize and cause leakage of the inner membrane of colistin-resistant *K. pneumoniae* MKP103. Depolarization is caused by many host defense peptides including those that do not form pores [73, 74]. However, predominantly pore forming peptides can cause leakage of the cytoplasmic components [75, 76]. With the lesser degree of outer membrane disruption observed with our more active peptide variants, we hypothesize they are penetrating to the inner membrane but their mechanism of interaction with the inner membrane is variable depending on the mutations introduced. We evaluated changes in membrane polarity across the inner membrane with 3,3’-dipropylthiadicarbocyanine Iodide (DiSC3) [77]. This cationic dye inserts into polarized inner membranes quenching its fluorescence and depolarization of the inner membrane results in release of the dye and an increase in fluorescence. When testing the single mutation variants (R9P, R10L and R11P) at 31 µmol L^-1^, only our beta sheet single peptide variant (R9P) depolarized the inner membrane of resistant *K. pneumoniae* MKP103 to the extent of the parent protegrin-1, while our more active beta turn variant (R11P) depolarized the inner membrane less than our parent peptide **(Figure 6A)**. Our less active beta turn variant (R10L) comparatively exhibited the least amount of depolarization toward MKP103. When testing the double mutation peptide variants (R9P, R10L and R10L, 11P) at 31 µmol L^-1^, both variants depolarized the inner membrane less than the parent protegrin-1 **(Figure 6B)**. Overall, all peptide variants depolarized the inner membrane greater than the polymyxin B, tested at 31 µmol L^-1^. We continued our investigation of changes in inner membrane polarity with our *ΔphoP* transposon mutant. We found that unlike our parental isolate, both our R9P and R11P depolarize the inner membrane to the same extent as parent protegrin-1 **(Figure 6C)**. Our beta turn single variant (R10L) does continue to have the lowest depolarization ability of all our peptide variants. This trend continues with our double variants, where both our beta turn and beta sheet double variants exhibit depolarization on par with our parent peptide **(Figure 6D)** rather than lower depolarization, as seen in our parent MKP103.

**Figure 6.**
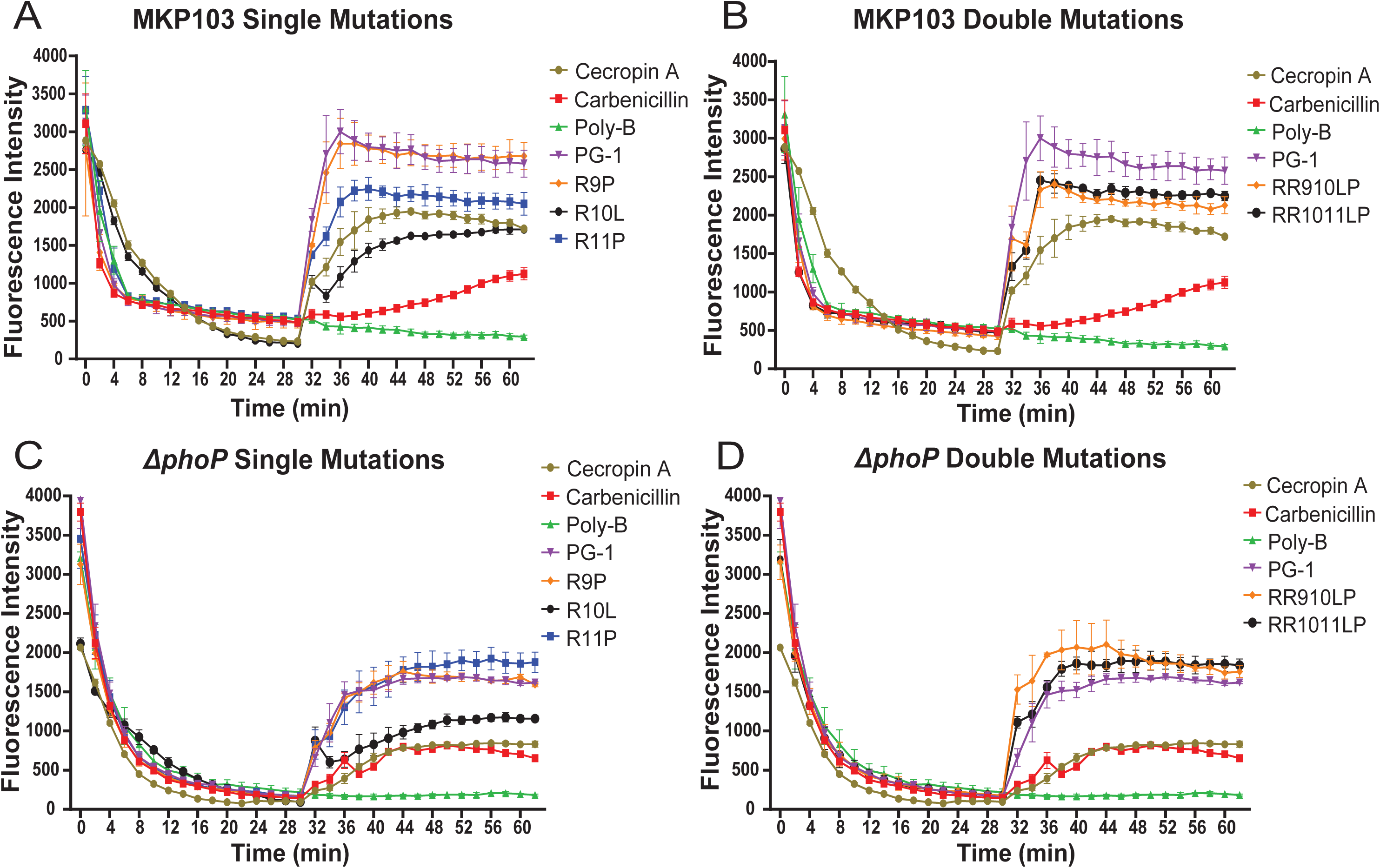
The arginine to proline mutation in the beta-sheet transition shows increased membrane depolarization. Membrane depolarization shows less active single peptide variant (R9P) depolarizing more effectively than our more active single peptide variant (R11P). *K. pneumoniae* MKP103 tested with single peptide variants (A) and double peptide variants (B). *ΔphoP* transposon mutant tested against single peptide variants (C) and double peptide variants (D). Peptides were tested at a concentration of 31 µmol L^-1^ with error shown as ±SEM.

To assess for changes in cytoplasmic leakage, we utilized ortho-nitrophenyl-β-galactoside (ONPG), a substrate that is hydrolyzed to ortho-nitrophenol (ONP) by cytoplasmic β-galactosidase [78]. When testing the single mutation peptide variants (R9P and R11P) at 31 µmol L^-1^ , we found although both variants resulted in leakage similar to the parental protegrin-1, there was a delay in the time to cause equal leakage (**Figure 7A**). Specifically, the R11P peptide variant matched the ONP levels induced by the parental protegrin-1 after 25 minutes, while the R9P peptide variant did not match the leakage until 40 minutes. Surprisingly, our beta turn single variant (R10L) resulted in very little inner membrane leakage, displaying levels less than our inactive polymyxin B. When testing the double mutation variants (R9P,R10L and R10L,R11P) at 31 µmol L^-1^, our beta sheet double peptide variant resulted in a decrease in the ONP levels compared to the parental protegrin-1 and displayed levels similar to the inactive polymyxin B (**Figure 7B**). The beta turn double peptide variant displayed a delay in fluorescence compared to the parental protegrin-1 at this concentration **(Figure 7B)**. Investigation with our *ΔphoP* transposon mutant resulted in an overall inability to effectively disrupt the inner membrane with all peptide variants **(Figure 7C and D)**. We also find that all peptide variants behave similarly to parent protegrin-1 regardless of differences in antimicrobial activity. Our beta turn single variant (R10L), however, still behaves similarly to our inactive polymyxin B. **(Figure 7C)**. Collectively, our results show that all peptide variants depolarize the inner membrane but changes in cytoplasmic leakage correlate best with the peptide variants MICs.

**Figure 7.**
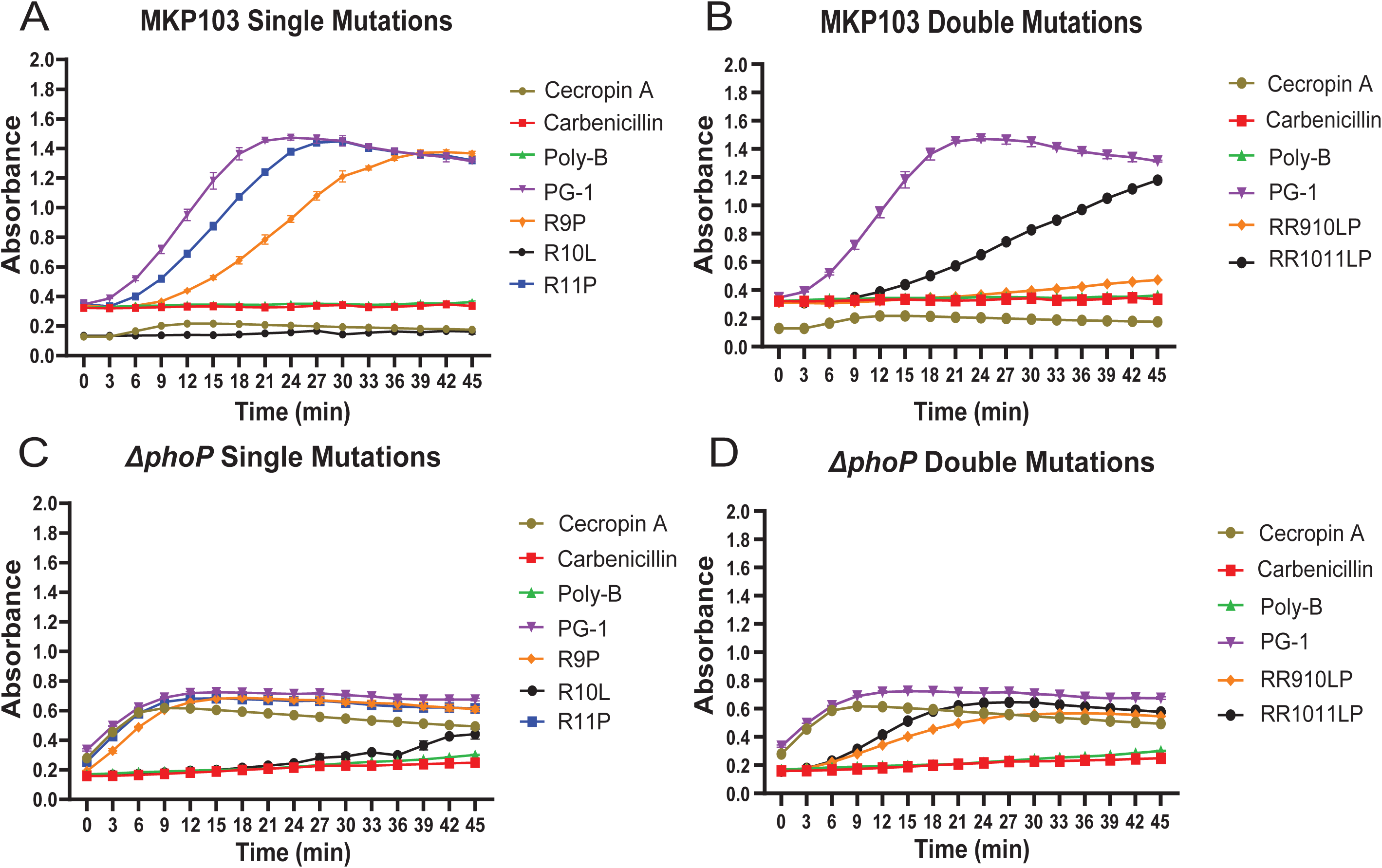
The arginine to proline mutation in the beta-sheet transition limits cytosolic leakage in single and double mutation variants. Inner membrane leakage shows more active single peptide variant (R11P) disrupting more effectively than our less active single peptide variants (R9P, R10L). *K. pneumoniae* MKP103 tested with single peptide variants (A) and double peptide variants (B). *ΔphoP* transposon mutant tested against single peptide variants (C) and double peptide variants (D). Peptides were tested at a concentration of 31 µmol L^-1^ with error shown as ±SEM.

### Molecular modeling shows octameric pore collapse and loss of beta sheet character in beta sheet peptide variants

We found the membrane leakage to correspond to the decrease in antimicrobial activity observed with our peptide variants suggesting a deficiency in the pore formation potential of the peptide variants. Alpha helical peptides like ll-37 have been shown to dimerize before insertion into the membrane [79]. However, protegrin-1 has been shown to dimerize within the phospholipid bilayer of the inner membrane ultimately leading to octameric pore formation [55]. Therefore, to provide a full spectrum of the potential of the mutations in the peptide variants we modeled the peptide variants before and after interaction with the bacterial membrane, as well as their octameric pore conformations when fully inserted. We also chose to eliminate our beta turn single peptide variant (R10L) due to its inability to cause any inner membrane leakage.

We modeled our octameric pores utilizing AlphaFold2 with the sequences aligned in the N- to C-termini direction. Images were visualized and diameter distances generated utilizing ChimeraX. Compared to our parent peptide **(Figure 8A)**, our less active single peptide variant (R9P) **(Figure 8B)** exhibits a reduction in diameter in our octameric pore model (∼20Å to 11Å, respectively). In contrast, our more active single peptide variant (R11P) **(Figure 8C)** changes the orientation of the hypothetical pore but maintains pore diameter at ∼20Å. When comparing our parent peptide to our double peptide variants we find our beta sheet double peptide variant (R9P, R10L) also exhibiting a pore reduction **(Figure 8D)** to ∼14-15 Å, and our beta turn double peptide variant (R10L, R11P) **(Figure 8E)** maintained the pore diameter (∼ 18Å). This suggests that perhaps the monomeric structures involved in pore formation are differentially impacted by our introduced mutations.

**Figure 8.**
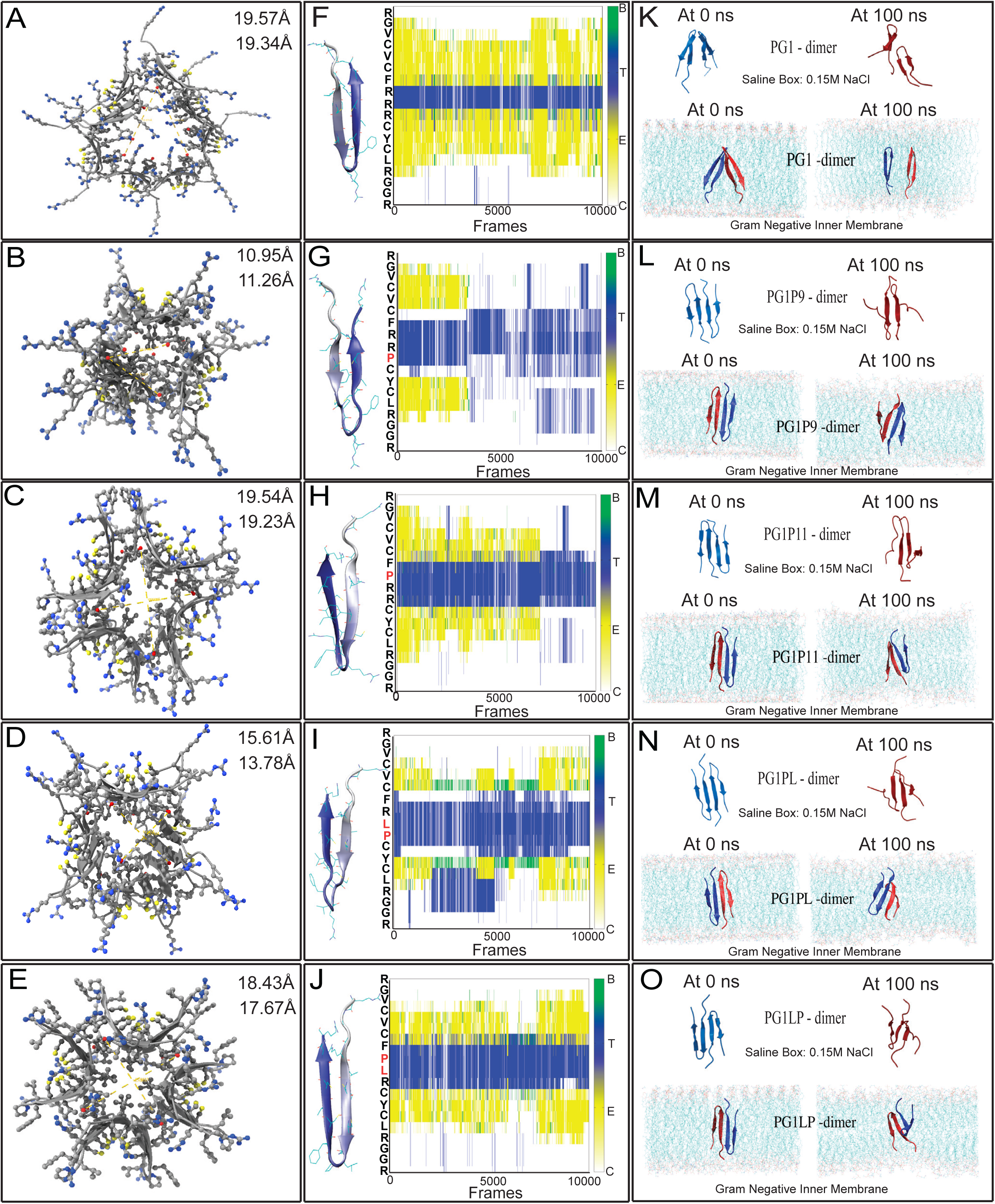
Molecular modeling shows octameric pore collapse and loss of beta sheet character in beta sheet peptide variants. Peptide variants are shown modeled as octameric pores in Figure 8A**-8E** using AlphaFold2 and visualized using ChimeraX for parent protegrin-1 (**A**), R9P (**B**), R11P (**C**), RR910PL (**D**) and RR1011LP (**E**). Amino acid sequences were aligned in the N to C termini direction. Figures 8F**-8J** show monomeric peptide variants simulated using GROMACs and observed for changes in secondary structure parent protegrin-1 (**F**), R9P (**G**), R11P (b), R9P, R10L (**I**) and R10L, R11P (**J**). The heat map labels show the color coding for B = Beta Bridge, T =Turn, E = Extended Conformation, C = Coil peptide structural components. Figures 8K**-8O** show dimeric peptide variants simulated using GROMACs in both a salt box (0.15M NaCl at 310K) (top) and a general Gram-negative inner membrane (bottom). Snapshots are shown for parent protegrin-1 (**K**), R9P (**L**), R11P (**M**), R9P, R10L (**N**) and R10L, R11P (**O**) at 0 and 100ns for both simulated environments.

To better understand the peptide variants behavior within the octameric pore, we modeled our peptide as monomeric subunits and observed how the structure of the peptide became altered in solution. To do this we utilized GROMACS, a molecular dynamics software that uses different algorithms to model the behavior of the peptide and its interactions with individual molecules over a span of time. These mutations were modeled in an environment of 0.15M NaCl at 310K over the course of 100ns to observe changes in the structural conformation of each peptide mutant [80]. We use Secondary Structure Predictors (DSSP) to observe changes in beta-sheet character over the simulation period. Compared to the parent peptide **(Figure 8F)**, the proline mutation at position 9 resulted in a shortened beta-sheet in its initial structure and an early loss of beta sheet character, at approximately 4000 frames (40 ns) that was never regained throughout the simulation **(Figure 8G)**. However, at position 11, the initial structure remained close to that of the parent peptide and the loss of beta character occurred after the halfway point of the simulation, at approximately 60000 frames (60 ns) (**Figure 8H).** When assessing the double peptide variants, the beta sheet region variant (R9P, R10L) does display a minor level of beta-sheet character throughout the simulation, however it is interwoven with regions of disorder. Furthermore, its initial structure does have shorter beta sheets when compared to the parent peptide **(Figure 8F and 8I).** Finally, our beta turn double peptide variant, (R10L, R11P) results in no changes to its initial structure when compared to the parent peptide. This double mutation fully loses its beta sheet character at approximately 60000 frames (60 ns) and regains it at approximately 80000 frames (80 ns), which may account for its ability to form a correctly oriented pore **(Figure 8E and 8J)**.

Protegrin-1 dimerizes before aggregating into an octameric pore inside the inner membrane. To better assess how our mutations altered peptide structure we assessed changes in its ability to form a dimer. Once again, we utilized GROMACs and an environment of both 0.15M NaCl at 310K and a general Gram-Negative inner membrane, over the course of 100ns [80, 81]. Before and after snapshots were taken to observe changes in the secondary structure of each mutated dimer compared to the parent **(Figure 8K)**. Parent protegrin-1 does not seem to align itself as a dimer in the salt box at the beginning or end of the simulation. This is also true in the Gram-Negative inner membrane simulation where it overlaps at the beta turn region at the beginning and orients across from itself after 100ns. This is unlike both single peptide variants **(Figure 8L and 8M)**. Both align themselves together at the beginning of the simulation.

Our R9P only aligns one beta sheet at the end of the simulation, however, it does display a lengthened beta-sheet. This is unlike R11P, which does not display a lengthened beta sheet at the end of the simulation. Inside the Gram-Negative environment, both single mutant variants orient very similarly at the beginning of the simulation, however at the end they orient themselves diagonally in opposing directions. Our beta sheet double peptide variant found in the screen (R9P, R10L) also aligns with itself at the beginning of the simulation and displays some alignment towards the end **(Figure 8N)**. Our other beta turn double peptide variant, R10L, R11P also aligned itself at the beginning of the simulation **(Figure 8O)**. Unlike all the other mutations, however, there is no alignment at the end of the simulation. In the Gram-negative inner membrane model, both double peptide variants exhibit diagonal alignment of both of their beta sheets, albeit in opposing directions. Looking at how each dimer behaves over the course of the simulation, protegrin-1 has a larger and more variable center of mass than all our point mutations, suggesting a less stable conformation **(Figure S5A)**. Our mutated peptide variants all behaved similarly, having a lower center of mass compared to the parent peptide. This suggests less movement and could explain why there was partial alignment of the peptide variants in the salt environment after 100ns **(Figure 8K-O)**. Furthermore, there were fewer contacts that existed between the dimers of parent protegrin-1 over the course of the salt box simulation **(Figure S5B)**. Meaning this set of monomers were not interacting with each other, explaining why there is no alignment at the end of the simulation **(Figure 8K-O).** Aside from our RR910PL point mutations, all other mutations have a larger and about the same number of contacts across the simulations. Our RR910PL mutation had two instances of a higher number of contacts, from approximately 4000 to 6000 ns, and 8000 to 8500 ns. This change in the number of contacts may explain its change in alignment compared to our other point mutations. Within the context of our Gram-negative inner membrane simulation our parent peptide also exhibited a large center of mass compared to our peptide variants, which could explain why at the end of the simulation they look as if on opposing sides of a potential pore **(Figure 8K, S5C)**. Furthermore, there is a decrease in the number of contacts between each chain over the course of the simulation **(Figure S5D)**. However, unlike in our salt box there is a much larger gap in the initial number of contacts. This could be due to how they initially intercalated within the inner membrane bilayer. Taken together these hypothetical models show our peptide variants with beta sheet (R9P and R9P, R10L) mutations impact pore formation and our monomeric subunits exhibit a rapid loss of beta sheet character not seen with our beta turn variants (R11P and R10L, R11P).

## Discussion

Carbapenem *K. pneumoniae* resistant isolates are now acquiring resistance to the last resort antibiotic known as polymyxin E or colistin [7, 17–20]. This resistance is attributed to modifications to the LPS by the PhoPQ two component system [26–29]. Here we corroborate previous literature stating that host-defense peptides are not equally affected by these LPS modifications as colistin [30–32,37,38], especially protegrin-1 **(Figure 2B)**. The absence of resistance, combined with its well characterized structure and robust antimicrobial activity [54–56] made protegrin-1 an ideal candidate to investigate colistin resistance in *K. pneumoniae*.

Utilizing our dmSLAY genetic screening platform in combination with quantitative next generation sequencing we found five amino acid mutations that were abundant in the population of plasmids retained following 4-hour induction, indicating they may alter the antimicrobial activity of protegrin-1. First, an arginine substituting a glycine (G2R) in the tail region, potentially altering charge localization of the peptide to the point where it no longer retained activity. An isoleucine substituting a leucine (L5I) in the first beta-sheet potentially adding more bulk to the peptide. A proline in the triple arginine motif within the beta-sheet to turn transition (R9P) that could introduce a kink and potentially disrupting the function of the peptide. Finally, a leucine in both the turn region (R10L) and second beta-sheet (F12L) that would allow for less binding of the peptide when compared to an arginine or phenylalanine, respectively. We also found that our beta sheet proline and beta turn leucine mutations (R9P, R10L) most commonly occurred together in peptide variants retained within the population. To test the effects of these mutations independently without other peptide modifications, we directly cloned them into our surface display genetic platform and performed growth analysis, revealing the beta sheet region arginine to proline (R9P) and beta turn region arginine to leucine (R10L) mutations has the greatest impact on the antimicrobial activity of protegrin-1, and combining these mutations (R9P,R10L) resulted in a complete loss of antimicrobial activity **(Figure 3D)**. Overall, we were surprised that more mutations resulted in possible structural changes rather than changes to charge in the population retained in the population in the dmSLAY analysis, because charge has been the major factor shown to contribute to bypassing the Gram-negative membrane [82]. This included those that added arginine residues to the unstructured tail region (G2R and G3R) increasing the charge of the peptide but not antimicrobial activity.

We then synthesized peptide variants to validate the surface display system and test our hypothesis that structure potentially supersedes charge. Specifically, we synthesized R11P and R10L,R11P peptide variants that placed the proline residue in the turn region of the triple arginine motif rather than the beta sheet transition region. We found then when testing these variants against MKP103 that our R11P peptide variant retains antimicrobial activity and our R9P peptide variant loses activity, despite a charge reduction on +1 in both peptide variants **(Table 4)**. Furthermore, when comparing the activity of our peptide variants against MKP103 and transposon mutants of genes under the regulation of PhoP, we find the fold change difference in activity is lower with our variants in the beta turn region versus the beta sheet region **(Figure 4B)**. These modifications both resulted in a loss of charge but did not equally lose antimicrobial activity, challenging the paradigm that charge is not necessarily most important aspect in the ability of host defense peptide ability to overcome colistin resistance. To further test this hypothesis, we tested the addition of alanine and lysine modifications to positions 9 and 11 on protegrin-1. Surprisingly, we found these smaller amino acids when substituted at position 11 (R11A and R11K) resulted in a less antimicrobial activity towards colistin-resistant *K. pneumoniae* and this activity is rescued when testing mutants of colistin resistance modifying genes. These findings reinforce our hypothesis that small changes to structure, not just the loss of the cystine residues can impact antimicrobial acidity of protegrin-1 more than a +1-charge reduction.

To understand the changes in activity observed with the synthesized peptide variants, we investigated their interactions with the outer and inner bacterial membrane. Outer membrane permeability accounted for some of our activity changes as both our double variants exhibited higher fluorescence compared to the parent peptide, suggesting they are getting stuck within the outer membrane **(Figure 5A)**. This is in line with what has been shown with polymyxin B nonapeptide lacking the lipid tail that can still increase penetration through the outer membrane without inner membrane lysis [83]. The differences in antimicrobial activity between our single proline peptide variants (R9P and R11P) correlated will with inner membrane permeability. We found R9P peptide variant exhibited a slower accumulation of inner membrane leakage compared to R11P, which behaved similarly to the parent peptide **(Figure 7A),** revealing changes in pore formation ability may be the reason for the decrease in antimicrobial activity with the R9P peptide variant. These findings reveal the small changes to protegrin-1 structure, not just the major changes observed with cysteine bond removal [62], can impact inner membrane leakage of the colistin-resistant *K. pneumoniae* membrane.

At the inner membrane protegrin-1 forms octameric pores, which cause cytoplasmic leakage [55].To understand why our peptide variants behaved differently at the inner membrane we modeled our variants as octameric pores. The resulting hypothetical models show a reduction in pore diameter and visual pore collapse in both beta sheet peptide variants (R9P and R9P, R10L) **(Figure 8B and 8D)** but not with our beta turn peptide variants (R11P and R10L, R11P) **(Figure 8C and 8E)**, where we found they maintain pore integrity and diameter, only changing in peptide orientation within the pore. To understand the structural instabilities leading to decreased pore formation we used GROMACs to observe changes in the secondary structure in physiological salt conditions and within a lipid environment. When assessing monomeric structural changes within salt conditions, we found the R9P beta sheet peptide variant loses beta sheet structure early in the simulation while the R11P beta turn variant can maintain beta sheet structure halfway through the simulation **(Figure 8G and 8H)**. The R9P, R10L beta sheet peptide variant exhibits a much lower beta sheet character throughout the entirety of the simulation **(Figure 8I)**, while the R10L, R11P beta turn peptide variant behaves much like R11P peptide variant **(Figure 8H and 8J)**. These differences in monomeric behavior allowed us to hypothesize that perhaps pore formation is hindered depending on at what point a variant loses its beta sheet character. When assessing the dimer formation and stability in both salt and lipid environments we found overall the orientation of the dimer changes with all variants, where the parental peptide dimerizes in parallel orientation and the variants in anti-parallel orientation. In all the peptides maintained structural integrity in the lipid environment more than the salt environment even at the beginning of the simulation (0 seconds) (**Figure 8K-8O**). Furthermore, we found only the parental protegrin-1 dimer separated after 100 ns (**Figure 8K**). These factors suggest strong dimerization may not be as important for pore formation with protegrin-1.

In conclusion, colistin and host defense peptides antimicrobial activity has been largely attributed to charge, as peptides with a higher positive charge will be more attracted to the negatively charged membrane. We show that with colistin-resistant *K. pneumoniae*, small changes in structure can impact antimicrobial activity more than decreasing the peptide charge from +6 to +5. This work represents a step towards understanding how to prevent activity loss due to colistin resistance membrane modifications when designing therapeutics for *K. pneumoniae*.

## Acknowledgements

This work is supported by the National Institutes of Health R00AI163295 to RMF; Cesar de la Fuente-Nunez holds a Presidential Professorship at the University of Pennsylvania and acknowledges funding from the Procter & Gamble Company, United Therapeutics, a BBRF Young Investigator Grant, the Nemirovsky Prize, Penn Health-Tech Accelerator Award, and the Dean’s Innovation Fund from the Perelman School of Medicine at the University of Pennsylvania. Research reported in this publication was supported by the Langer Prize (AIChE Foundation), the National Institute of General Medical Sciences of the National Institutes of Health under award number R35GM138201, and the Defense Threat Reduction Agency (DTRA; HDTRA11810041, HDTRA1-21-1-0014, and HDTRA1-23-1-0001). We would like to thank Bryan Davies for his support and guidance on this work and Cory DuPais for teaching Christina how to process the SLAY library from the Illumina sequencing reads.

## Author Contributions

R.M.F. conceived the idea and designed the experimentation. Initial library creation performed by N.R, J.R, and R.M.F. Library analysis and direct cloning experiments performed by C.D. Antimicrobial activity assessments and membrane analyses performed by S.A. and C.D. Molecular Dynamic simulations performed by C.d.l.F.-N. and R.K. Biorender and AlphaFold images produced by C.D. Text was written by R.M.F and C.D, and edits were discussed with C.d.l.F.-N., R.K. and J.R.

## Materials and Methods

### Bacterial strains and peptides

All bacterial isolates used in this study are listed in **Supplemental Table S1**. The peptides used in this study are listed in **Table 1** with their molecular weight, length, charge, and secondary structure. Pretegrin-1 was ordered through Anaspec for the comparison to other host defense peptides in **Figure 2**. Arenicin and Bac7(1-35) were purchased from Novopro because they were unavailable through Anaspec. For consistency with our synthesized variants, protegrin-1 parent and peptide variants were ordered from Novopro at >90% purity (https://www.novoprolabs.com) for comparing antimicrobial activity of amino acid substitutions. All peptides were resuspended in ultra-purified water at 10 mg mL^-1^ and stored at −20 °C.

### Minimal inhibitory concentration assays

All minimal inhibitory concentration assays were performed using the broth dilution method in 96-well plates with the peptides tested by serial diluting in 0.2% BSA 0.01% acetic acid solution [84]. Bacteria were synchronized to mid-log phase and standardized to a final optical density 600nm (OD600) of 0.001 in Mueller Hinton broth to yield ∼5×10^5^ CFU mL^-1^. The assay incubated for 20 hours at 37 °C before determining MICs. All MICs were performed in triplicate and repeated on separate days.

### RT-qPCR of *phoP* expression levels

Each *K. pneumoniae* strain was grown overnight in LB broth. The following day each strain was synchronized to exponential phase at 37°C for 3 hours. RNA was extracted after bacteria had been synchronized for 3 hours at 37°C. Following hot phenol chloroform extraction previously described [50] samples were resuspended in nuclease free H20 and quantified by Nanodrop. RNA were aliquoted and stored at −80 for RT-qPCR. Samples were reverse transcribed with SuperScript™ IV VILO™ Master Mix (11756050, Invitrogen) following the manufacturer’s instructions. RTqPCR was performed using PowerUp™ SYBR™ Green Master Mix (A25742, Applied Biosystems) following the manufacturer’s instructions, and the QuantStudio 3 Real-Time PCR System (A28567, Applied Biosystems). Primers were designed using Integrated DNA Technologies, Inc. (IDT)’s PrimerQuest tool (**Table S4**). Relative expression levels were calculated using the 2-ΔΔCt method [49] with *rpoD* as housekeeping control. Triplicate samples were used with error shown as ±SEM.

### Alpha Fold Images

Images were initially prepared using Google Collab and this link https://colab.research.google.com/github/sokrypton/ColabFold/blob/main/AlphaFold2.ipynb#scrollTo=11l8k--10q0C. Each sequence was input, and images were displayed using the ChimeraX visualizer. Primary and secondary structures were overlayed and mutated amino acids are shown in red.

### Ramachandran Plot Generation

Plots were generated by taking previously generated alpha fold files and displaying them with the visualizing tool from UCSF chimera. Program can be accessed by utilizing this link https://www.cgl.ucsf.edu/chimera/download.html. Plots were recolored and overlayed using Adobe Photoshop.

### dmSLAY protegrin-1 analysis

#### dmSLAY creations and screening

The protegrin-1 deep mutational library was created as previously described [60] and inserted into a tetracycline resistant pMMB67EH plasmid. The library was first electroporated into *E. coli* C3020 and >150,000 colonies were collected. The library was then transformed into *E. coli* SM10 λpir mating strain and >270,000 colonies were collected. Finally, conjugation was used to introduce the library to *K. pneumoniae* MKP103 and ∼200,000 colonies were collected. Samples were frozen in glycerol stocks until library screening. Triplicate samples *K. pneumoniae* MKP103 harboring the dmSLAY protegrin-1 library were thawed and back diluted to and OD600 0.1 and grown for one hour. The samples were then back diluted to OD600 0.01 in 5 mL terrific broth (tryptone, yeast, potassium phosphate dibasic, and glycerol) and triplicate samples were induced with 0.05 mM IPTG or uninduced (0mM IPTG) for three hours shaking (220rpm) at 37 °C followed by, bacterial population collection and subsequent plasmid extraction. Barcoded primers (Table S7) were used for PCR of the triplicate induced and uninduced plasmid populations for amplification of the 240 bp amplicon for Illumina sequencing. One lane of Illumina MiSeq run was used for amplicon coverage to allow for quantification and calculation of log2 fold change between the induced and uninduced samples.

#### Bioinformatic Analysis

The next generation sequencing reads were process by having their forward and reverse sequences merged together and adaptor sequences trimmed. For this we used a unique modification of DESQ2, a package associated with R Studio, typically used to investigate the presence of genes within a population. More specifically, we merged forward and reverse raw sequences from a previously made protegrin-1 library, and adaptors were trimmed from both 5’ and 3’ end of the sequence. Reads were counted and those occurring less than 50 times within the population were removed from analysis to allow for the possibility of machine error. Translated DNA sequences were then compared to the reads of the uninduced and induced populations using the Deseq2 package in R studio. To allow for analysis of specifically those reads that did not kill the bacteria we eliminated reads with a log 2-fold change below 0. From this data each amino acid mutation at each position in the total sequence variants were counted to quantify their occurrence within the population of peptide variants.

### Direct Cloning of Peptide Variants

Procedure adapted from previous methods [59]. Briefly, protegrin-1 parent plasmid was used as a template for cloning via restriction enzymes on the broad host plasmid PMMB67EH with a tetracycline resistance cassette. Sequences were cloned with peptide variant primers listed in **Table S5** with homology to the tether sequence on the reverse primer. Restriction digestion was performed using KpmI and SalI cut sites. Each peptide variant was then transformed into *E.coli C320* (<u>NEB) electro</u>competent cells and sequence variants were confirmed by sanger sequencing. Plasmid was extracted from *E. coli* using Qiagen mini-prep kit and electroporated into MKP103 for testing of antimicrobial activity with IPTG induction.

### Surface display validation of mutations identified in library screen

Growth curve procedure was adapted from previous methods [59]. Each *K. pneumoniae* strain with plasmids harboring peptide variants and control plasmids listed in **Table S6**. were grown overnight in LB broth and 1% glucose solution supplemented with 20µg/mL of tetracycline. The following day each strain was synchronized to exponential phase at 37°C for 3 hours. At that time the strains were standardized to an OD600 of 0.02. Bacterial strains were then added to culture tubes containing 2 mL LB, 0.5 mM IPTG, and 20 µg mL^-1^ tetracycline and grown shaking (220rpm) at 37°C over the course of 4 hours. At 0, 1, 2, and 4 hours 100 µL was removed from each culture tube and added to 900µL of 1XPBS for 1:10 serial dilutions that were then plated on LB agar with 50 µg/mL tetracycline. All experiments were plated in triplicate and incubated overnight at 37°C, and colony forming units per mL (CFU mL^-1^) was calculated with error reported as ±SEM.

### Outer Membrane Permeability

Procedure adapted from previous methods [69–71] Each *K. pneumoniae* strain was grown overnight in LB broth. The following day each strain was synchronized to exponential phase at 37°C for 3 hours. The cells were harvested by centrifugation (3434 rcf for 10 min) washed twice with buffer A (5mM HEPES, 5mM glucose, pH 7.2) and resuspended in buffer A to an OD600 = 0.1. Then 100 μL Buffer A with 20 μM NPN was added to 1000 μL washed cells in a clear-bottom black walled 96-well plate. Peptide was added to a final concentration of 64 μg/mL and fluorescence was monitored at an excitation of 350nm and emission of 420nm for 30 minutes at 3-minute intervals. The read form the no cell control wells were removed from each treated well to normalize for background. All assays were performed in triplicate with error reported as ±SEM.

### Membrane Depolarization

The procedure used was adapted from a previous method [77]. Briefly, each *K. pneumoniae* strain was grown overnight in LB broth. The following day each strain was synchronized to exponential phase at 37°C for 3 hours. The cells were harvested by centrifugation (3434 rcf <u>for 10 min) washed twice with</u> buffer A (5mM HEPES, 5mM glucose, pH 7.2) and resuspended in buffer A and 100mM KCl to an OD600 = 0.1, along with 20mM DiSC3 dye. Bacteria was added to clear-bottom black walled 96-well plate and standardized for 30 minutes monitoring the florescence changes using a Biotech Synergy H1 plate reader with fluorescence kinetic reads set with excitation of 622nm and emission of 670nm for 30 minutes at 2-minute intervals. Peptide was then added to a final concentration of 64 μg mL^-1^ and fluorescence changes were monitored for 30 minutes at 2-minute intervals using excitation of 622nm and emission of 670nm. All assays were performed in triplicate with error reported as ±SEM.

### Inner Membrane Leakage

The procedure used was adapted from a previous method [77, 78]. Each *K. pneumoniae* strain was grown overnight in LB broth. The following day each strain was synchronized to exponential phase at 37°C for 3 hours. The cells were harvested by centrifugation (3434 rcf for 10 min) washed once with 0.1M sodium phosphate buffer (26mM NaH2PO4, 77mM Na2HPO4, pH 7.4) and resuspended in 0.1M sodium phosphate buffer to an OD600 = 0.1 along with 10mM ONPG. The peptide were then serially diluted in sodium phosphate buffer in a clear-bottom walled black 96-well plate, bacteria were added, and absorbance was read using a Biotech Synergy H1 plate reader at 410nm for 45 minutes with reads set at 3-minute intervals.

### Molecular Dynamics Simulations

GROMACs version 2025.0 located https://www.gromacs.org/index.html was used to perform the analysis of the peptide variant monomers as well as dimers in both a saline medium (ionic strength: 0.15 M NaCl) and a general Gram-negative inner membrane environment located https://charmm-gui.org/?doc=archive&lib=biomembrane. This analysis analyzed changes in peptide secondary structure as well as fluctuations in center of mass and residue contacts with peptide variant dimers. Snapshots of the peptide monomers were taken at 0ns, and peptide dimers were taken at 0 and 100ns to all for visualization of structural changes in each subunit.

## References

1. Paczosa, M.K. and J. Mecsas, Klebsiella pneumoniae: Going on the Offense with a Strong Defense. Microbiol Mol Biol Rev, 2016. 80(3): p. 629–61.

2. Asokan, S., et al., Klebsiella pneumoniae: A growing threat in the era of antimicrobial resistance. The Microbe, 2025. 7: p. 100333.

3. Navon-Venezia, S., K. Kondratyeva, and A. Carattoli, Klebsiella pneumoniae: a major worldwide source and shuttle for antibiotic resistance. FEMS Microbiol Rev, 2017. 41(3): p. 252–275.

4. Shah, A.A., et al., Emerging challenges in Klebsiella pneumoniae: Antimicrobial resistance and novel approach. Microbial Pathogenesis, 2025. 202: p. 107399.

5. Reynolds, D., et al., The threat of multidrug-resistant/extensively drug-resistant Gram-negative respiratory infections: another pandemic. Eur Respir Rev, 2022. 31(166).

6. Antimicrobial Resistance, C., Global burden of bacterial antimicrobial resistance in 2019: a systematic analysis. Lancet, 2022. 399(10325): p. 629–655.

7. Petrosillo, N., F. Taglietti, and G. Granata, Treatment Options for Colistin Resistant Klebsiella pneumoniae: Present and Future. J Clin Med, 2019. 8(7).

8. Wyres, K.L., M.M.C. Lam, and K.E. Holt, Population genomics of Klebsiella pneumoniae. Nat Rev Microbiol, 2020. 18(6): p. 344–359.

9. Sati, H., et al., The WHO Bacterial Priority Pathogens List 2024: a prioritisation study to guide research, development, and public health strategies against antimicrobial resistance. Lancet Infect Dis, 2025.

10. Mondal, A.H., et al., A Review on Colistin Resistance: An Antibiotic of Last Resort. Microorganisms, 2024. 12(4).

11. Rojas, L.J., et al., Colistin Resistance in Carbapenem-Resistant Klebsiella pneumoniae: Laboratory Detection and Impact on Mortality. Clin Infect Dis, 2017. 64(6): p. 711–718.

12. van Duin, D. and Y. Doi, Outbreak of Colistin-Resistant, Carbapenemase-Producing Klebsiella pneumoniae: Are We at the End of the Road? J Clin Microbiol, 2015. 53(10): p. 3116–7.

13. Luterbach, C.L., et al., Transmission of Carbapenem-Resistant Klebsiella pneumoniae in US Hospitals. Clin Infect Dis, 2023. 76(2): p. 229–237.

14. Horvath, M., et al., Virulence Characteristics and Molecular Typing of Carbapenem-Resistant ST15 Klebsiella pneumoniae Clinical Isolates, Possessing the K24 Capsular Type. Antibiotics (Basel), 2023. 12(3).

15. Marsh, J.W., et al., Evolution of Outbreak-Causing Carbapenem-Resistant Klebsiella pneumoniae ST258 at a Tertiary Care Hospital over 8 Years. mBio, 2019. 10(5).

16. Chen, L., et al., Epidemic Klebsiella pneumoniae ST258 Is a Hybrid Strain. mBio, 2014. 5(3): p. 10.1128/mbio.01355-14.

17. Bogdanovich, T., et al., *Colistin-resistant,* Klebsiella pneumoniae carbapenemase (KPC)-producing Klebsiella pneumoniae belonging to the international epidemic clone ST258. Clin Infect Dis, 2011. 53(4): p. 373–6.

18. Lapp, Z., et al., Fitness barriers to spread of colistin resistance overcome by first establishing niche in patients with enhanced colistin exposure. medRxiv, 2021: p. 2021.06.11.21258758.

19. Pitt, M.E., et al., Octapeptin C4 and polymyxin resistance occur via distinct pathways in an epidemic XDR Klebsiella pneumoniae ST258 isolate. J Antimicrob Chemother, 2019. 74(3): p. 582–593.

20. Arena, F., et al., Colistin Resistance Caused by Inactivation of the MgrB Regulator Is Not Associated with Decreased Virulence of Sequence Type 258 KPC Carbapenemase-Producing Klebsiella pneumoniae. Antimicrob Agents Chemother, 2016. 60(4): p. 2509–12.

21. Diani, E., et al., Colistin: Lights and Shadows of an Older Antibiotic. Molecules, 2024. 29(13).

22. Khondker, A. and M.C. Rheinstädter, How do bacterial membranes resist polymyxin antibiotics? Communications Biology, 2020. 3(1): p. 77.

23. Hamel, M., J.M. Rolain, and S.A. Baron, The History of Colistin Resistance Mechanisms in Bacteria: Progress and Challenges. Microorganisms, 2021. 9(2).

24. Yu, Z., et al., Antibacterial mechanisms of polymyxin and bacterial resistance. Biomed Res Int, 2015. 2015: p. 679109.

25. Simpson, B.W. and M.S. Trent, Pushing the envelope: LPS modifications and their consequences. Nat Rev Microbiol, 2019. 17(7): p. 403–416.

26. Bray, A.S., et al., MgrB-Dependent Colistin Resistance in Klebsiella pneumoniae Is Associated with an Increase in Host-to-Host Transmission. mBio, 2022. 13(2): p. e03595–21.

27. Janssen, A.B., et al., Evolution of Colistin Resistance in the Klebsiella pneumoniae Complex Follows Multiple Evolutionary Trajectories with Variable Effects on Fitness and Virulence Characteristics. Antimicrob Agents Chemother, 2020. 65(1).

28. Li, L., et al., Roles of two-component regulatory systems in Klebsiella pneumoniae: Regulation of virulence, antibiotic resistance, and stress responses. Microbiol Res, 2023. 272: p. 127374.

29. Kim, S.J., et al., Roles of crrAB two-component regulatory system in Klebsiella pneumoniae: growth yield, survival in initial colistin treatment stage, and virulence. Int J Antimicrob Agents, 2024. 63(1): p. 107011.

30. Arroyo Luis, A., et al., The pmrCAB Operon Mediates Polymyxin Resistance in Acinetobacter baumannii ATCC 17978 and Clinical Isolates through Phosphoethanolamine Modification of Lipid A. Antimicrobial Agents and Chemotherapy, 2011. 55(8): p. 3743–3751.

31. Lee, K.K., et al., Heterogeneous efflux pump expression underpins phenotypic resistance to antimicrobial peptides. 2024, eLife Sciences Publications, Ltd.

32. MacNair, C.R., et al., Overcoming mcr-1 mediated colistin resistance with colistin in combination with other antibiotics. Nat Commun, 2018. 9(1): p. 458.

33. Brogden, K.A., Antimicrobial peptides: pore formers or metabolic inhibitors in bacteria? Nat Rev Microbiol, 2005. 3(3): p. 238–50.

34. Sengupta, D., et al., Toroidal pores formed by antimicrobial peptides show significant disorder. Biochimica et Biophysica Acta (BBA) - Biomembranes, 2008. 1778(10): p. 2308–2317.

35. Zhang, Q.Y., et al., Antimicrobial peptides: mechanism of action, activity and clinical potential. Mil Med Res, 2021. 8(1): p. 48.

36. Hancock, R.E. and A. Rozek, Role of membranes in the activities of antimicrobial cationic peptides. FEMS Microbiol Lett, 2002. 206(2): p. 143–9.

37. Fleeman, R.M. and B.W. Davies, Polyproline Peptide Aggregation with Klebsiella pneumoniae Extracellular Polysaccharides Exposes Biofilm Associated Bacteria. Microbiol Spectr, 2022. 10(2): p. e0202721.

38. Fleeman, R.M., et al., Defining principles that influence antimicrobial peptide activity against capsulated Klebsiella pneumoniae. Proc Natl Acad Sci U S A, 2020. 117(44): p. 27620–27626.

39. El-Sayed Ahmed, M.A.E., et al., Colistin and its role in the Era of antibiotic resistance: an extended review (2000-2019). Emerg Microbes Infect, 2020. 9(1): p. 868–885.

40. Lei, J., et al., The antimicrobial peptides and their potential clinical applications. Am J Transl Res, 2019. 11(7): p. 3919–3931.

41. Drayton, M., et al., Host Defense Peptides: Dual Antimicrobial and Immunomodulatory Action. Int J Mol Sci, 2021. 22(20).

42. Bader, M.W., et al., Recognition of Antimicrobial Peptides by a Bacterial Sensor Kinase. Cell, 2005. 122(3): p. 461–472.

43. Wu, K.-M., et al., Genome Sequencing and Comparative Analysis of Klebsiella pneumoniae NTUH-K2044, a Strain Causing Liver Abscess and Meningitis. Journal of Bacteriology, 2009. 191(14): p. 4492–4501.

44. Walker Kimberly, A., et al., A Klebsiella pneumoniae Regulatory Mutant Has Reduced Capsule Expression but Retains Hypermucoviscosity. mBio, 2019. 10(2): p. 10.1128/mbio.00089-19.

45. Rasheed, J.K., et al., Characterization of the Extended-Spectrum β-Lactamase Reference Strain, Klebsiella pneumoniae K6 (ATCC 700603), Which Produces the Novel Enzyme SHV-18. Antimicrobial Agents and Chemotherapy, 2000. 44(9): p. 2382–2388.

46. Cowan, S.T., et al., A classification of the Klebsiella group. J Gen Microbiol, 1960. 23: p. 601–12.

47. Ramage, B., et al., Comprehensive Arrayed Transposon Mutant Library of Klebsiella pneumoniae Outbreak Strain KPNIH1. J Bacteriol, 2017. 199(20).

48. Snitkin, E.S., et al., Tracking a hospital outbreak of carbapenem-resistant Klebsiella pneumoniae with whole-genome sequencing. Sci Transl Med, 2012. 4(148): p. 148ra116.

49. Livak, K.J. and T.D. Schmittgen, Analysis of relative gene expression data using real-time quantitative PCR and the 2(-Delta Delta C(T)) Method. Methods, 2001. 25(4): p. 402–8.

50. Aiba, H., S. Adhya, and B. de Crombrugghe, Evidence for two functional gal promoters in intact Escherichia coli cells. J Biol Chem, 1981. 256(22): p. 11905–10.

51. Mardirossian, M., et al., The Host Antimicrobial Peptide Bac71-35 Binds to Bacterial Ribosomal Proteins and Inhibits Protein Synthesis. Chemistry & Biology, 2014. 21(12): p. 1639–1647.

52. Barlow, D.J. and J.M. Thornton, Helix geometry in proteins. Journal of Molecular Biology, 1988. 201(3): p. 601–619.

53. Hauser, K., et al., Characterization of Biomolecular Helices and Their Complementarity Using Geometric Analysis. J Chem Inf Model, 2017. 57(4): p. 864–874.

54. Hollingsworth, S.A. and P.A. Karplus, A fresh look at the Ramachandran plot and the occurrence of standard structures in proteins. Biomol Concepts, 2010. 1(3-4): p. 271–283.

55. Bolintineanu, D.S., V. Vivcharuk, and Y.N. Kaznessis, Multiscale models of the antimicrobial peptide protegrin-1 on gram-negative bacteria membranes. Int J Mol Sci, 2012. 13(9): p. 11000–11011.

56. Fahrner, R.L., et al., Solution structure of protegrin-1, a broad-spectrum antimicrobial peptide from porcine leukocytes. Chem Biol, 1996. 3(7): p. 543–50.

57. Steinberg, D.A., et al., Protegrin-1: a broad-spectrum, rapidly microbicidal peptide with in vivo activity. Antimicrob Agents Chemother, 1997. 41(8): p. 1738–42.

58. Tachi, Y., S.G. Itoh, and H. Okumura, Molecular dynamics simulations of amyloid-beta peptides in heterogeneous environments. Biophys Physicobiol, 2022. 19: p. 1–18.

59. Tucker, A.T., et al., Discovery of Next-Generation Antimicrobials through Bacterial Self-Screening of Surface-Displayed Peptide Libraries. Cell, 2018. 172(3): p. 618–628 e13.

60. Randall, J.R., et al., *Deep mutational scanning and machine learning uncover antimicrobial peptide features driving membrane selectivity.* Res Sq, 2023.

61. Rodziewicz-Motowidlo, S., et al., Antimicrobial and conformational studies of the active and inactive analogues of the protegrin-1 peptide. FEBS J, 2010. 277(4): p. 1010–22.

62. Mohanram, H. and S. Bhattacharjya, Cysteine deleted protegrin-1 (CDP-1): Anti-bacterial activity, outer-membrane disruption and selectivity. Biochimica et Biophysica Acta (BBA) - General Subjects, 2014. 1840(10): p. 3006–3016.

63. Martin, M.J., et al., A panel of diverse Klebsiella pneumoniae clinical isolates for research and development. Microb Genom, 2023. 9(5).

64. Bishop, R.E., The lipid A palmitoyltransferase PagP: molecular mechanisms and role in bacterial pathogenesis. Mol Microbiol, 2005. 57(4): p. 900–12.

65. Brett, P.J., D. DeShazer, and D.E. Woods, Burkholderia thailandensis sp. nov., a Burkholderia pseudomallei-like species. Int J Syst Bacteriol, 1998. 48 **Pt** **1**: p. 317–20.

66. Jacobs Anna, C., et al., *AB*5075, a Highly Virulent Isolate of Acinetobacter baumannii, as a Model Strain for the Evaluation of Pathogenesis and Antimicrobial Treatments. mBio, 2014. 5(3): p. 10.1128/mbio.01076-14.

67. Jensen, K.F., The Escherichia coli K-12 “wild types” W3110 and MG1655 have an rph frameshift mutation that leads to pyrimidine starvation due to low pyrE expression levels. J Bacteriol, 1993. 175(11): p. 3401–7.

68. Minogue, T.D., et al., Complete Genome Assembly of Escherichia coli ATCC 25922, a Serotype O6 Reference Strain. Genome Announc, 2014. 2(5).

69. Loh, B., C. Grant, and R.E. Hancock, Use of the fluorescent probe 1-N-phenylnaphthylamine to study the interactions of aminoglycoside antibiotics with the outer membrane of Pseudomonas aeruginosa. Antimicrob Agents Chemother, 1984. 26(4): p. 546–51.

70. Muheim, C., et al., Increasing the permeability of Escherichia coli using MAC13243. Sci Rep, 2017. 7(1): p. 17629.

71. Richard, C.S.M., et al., Outer Membrane Integrity-Dependent Fluorescence of the Japanese Eel UnaG Protein in Live Escherichia coli Cells. Biosensors (Basel), 2023. 13(2).

72. Rizzetto, G., et al., Protegrin-1 and Analogues Against Acinetobacter baumannii: A Narrative Review. Pharmaceuticals (Basel), 2025. 18(3).

73. Huang, H.W., DAPTOMYCIN, its membrane-active mechanism vs. that of other antimicrobial peptides. Biochimica et Biophysica Acta (BBA) - Biomembranes, 2020. 1862(10): p. 183395.

74. Spindler, E.C., et al., Deciphering the mode of action of the synthetic antimicrobial peptide Bac8c. Antimicrob Agents Chemother, 2011. 55(4): p. 1706–16.

75. Mihaylova-Garnizova, R., et al., Antimicrobial Peptides Derived from Bacteria: Classification, Sources, and Mechanism of Action against Multidrug-Resistant Bacteria. Int J Mol Sci, 2024. 25(19).

76. Xiao, B., et al., Unlocking the Potential of Antimicrobial Peptides: Cutting-Edge Advances and Therapeutic Potential in Combating Bacterial Keratitis. Bioconjug Chem, 2025. 36(3): p. 311–331.

77. Moller, A.M., et al., Common and varied molecular responses of Escherichia coli to five different inhibitors of the lipopolysaccharide biosynthetic enzyme LpxC. J Biol Chem, 2024. 300(4): p. 107143.

78. Gravel, J., C. Paradis-Bleau, and A.R. Schmitzer, Adaptation of a bacterial membrane permeabilization assay for quantitative evaluation of benzalkonium chloride as a membrane-disrupting agent. MedChemComm, 2017. 8(7): p. 1408–1413.

79. Nielsen, J.E., et al., Self-Assembly of Antimicrobial Peptoids Impacts Their Biological Effects on ESKAPE Bacterial Pathogens. ACS Infect Dis, 2022. 8(3): p. 533–545.

80. Ferreira, A.F.L., et al., Defensins identified through molecular de-extinction. Cell Reports Physical Science, 2024. 5(9): p. 102193.

81. Pogozheva, I.D., et al., Comparative Molecular Dynamics Simulation Studies of Realistic Eukaryotic, Prokaryotic, and Archaeal Membranes. J Chem Inf Model, 2022. 62(4): p. 1036–1051.

82. Saxena, D., et al., Tackling the outer membrane: facilitating compound entry into Gram-negative bacterial pathogens. npj Antimicrobials and Resistance, 2023. 1(1): p. 17.

83. Trimble, M.J., et al., Polymyxin: Alternative Mechanisms of Action and Resistance. Cold Spring Harb Perspect Med, 2016. 6(10).

84. Wiegand, I., K. Hilpert, and R.E.W. Hancock, Agar and broth dilution methods to determine the minimal inhibitory concentration (MIC) of antimicrobial substances. Nature Protocols, 2008. 3(2): p. 163–175.

