## Supplemental File for "Defining the importance of the arginine loop region of protegrin-1 for antimicrobial activity towards colistin-resistant *Klebsiella pneumoniae*"

**Supporting Information**

**Figure 1. Fold Change MKP103 Colistin and Polymyxin B**

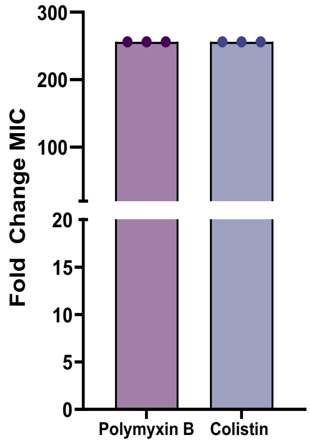


**Figure S1.** This figure shows the fold change difference in antimicrobial activity between wildtype MKP103 and its *ΔphoP* transposon mutant (KPNIH1_10030-701::T30). Colistin and polymyxin B were tested in triplicate at a maximum concentration of 31 μmol $L^{-1}$ (128 ug $mL^{-1}$).

**Figure 2. Possible Amino Acid Mutations with Mixed Base Primer**

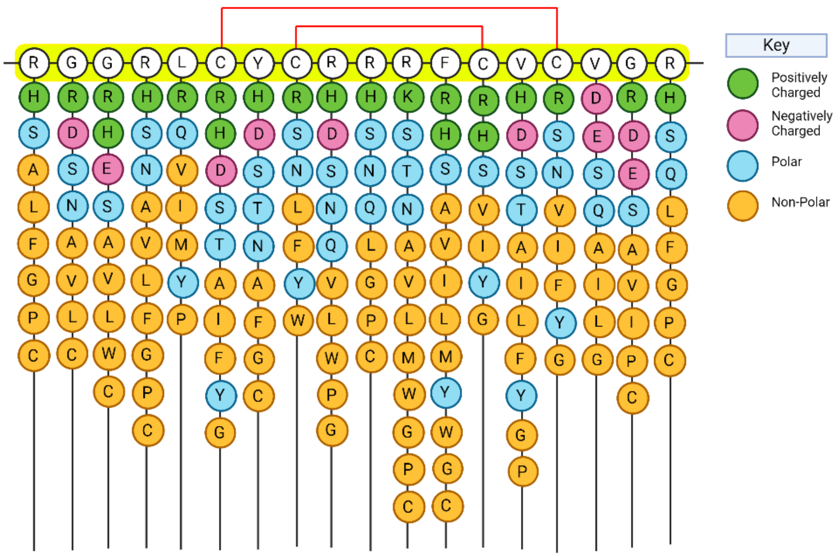


**Figure S2.** This figure is a schematic of our mixed base primer. The parent peptide sequence is denoted in white circles and highlighted in yellow. At each amino acid residue, colored circles indicate all potential amino acid mutations, and are color coded by amino acid class. Red lines at the top of the figure indicate the cysteines involved in each disulfide bridge in protegrin-1.

**Figure 3. Transposon Mutant Outer Membrane Permeability**

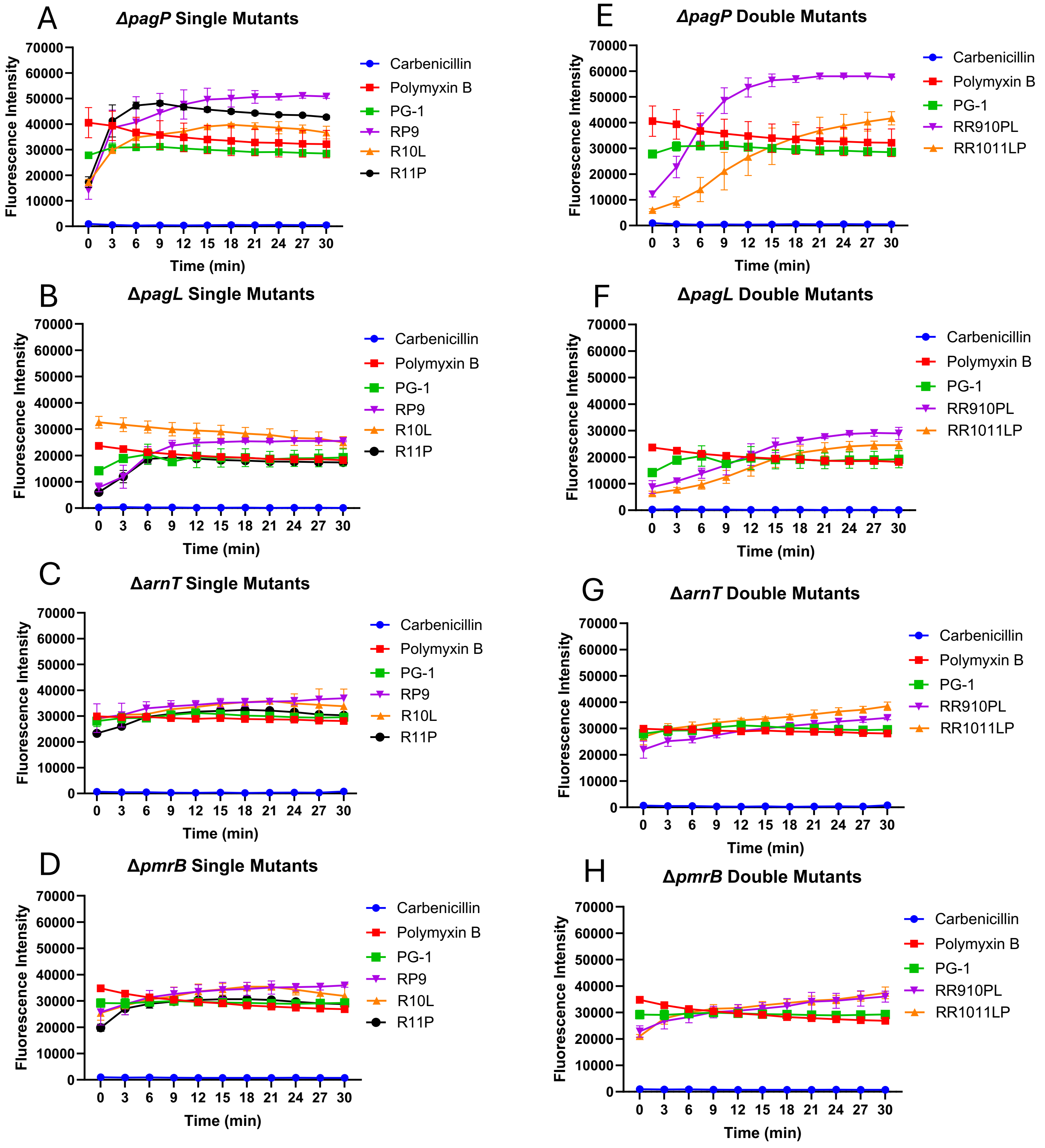


**Figure S3.** Transposon Mutant Outer Membrane Permeability shows variant changes in *ΔpagP.* Single mutant peptide variants assessed against *ΔpagP* transposon mutant (A) *ΔpagL* transposon mutant (B) *ΔarnT* transposon mutant (C) and *ΔpmrB* transposon mutant (D). Double mutant peptide variants assessed against *ΔpagP* transposon mutant (E) *ΔpagL* transposon mutant (F) *ΔarnT* transposon mutant (G) and *ΔpmrB* transposon mutant (H). All assays were performed in triplicate with error shown as ±SEM.

**Figure 4. Clinical Isolate Outer Membrane Permeability**

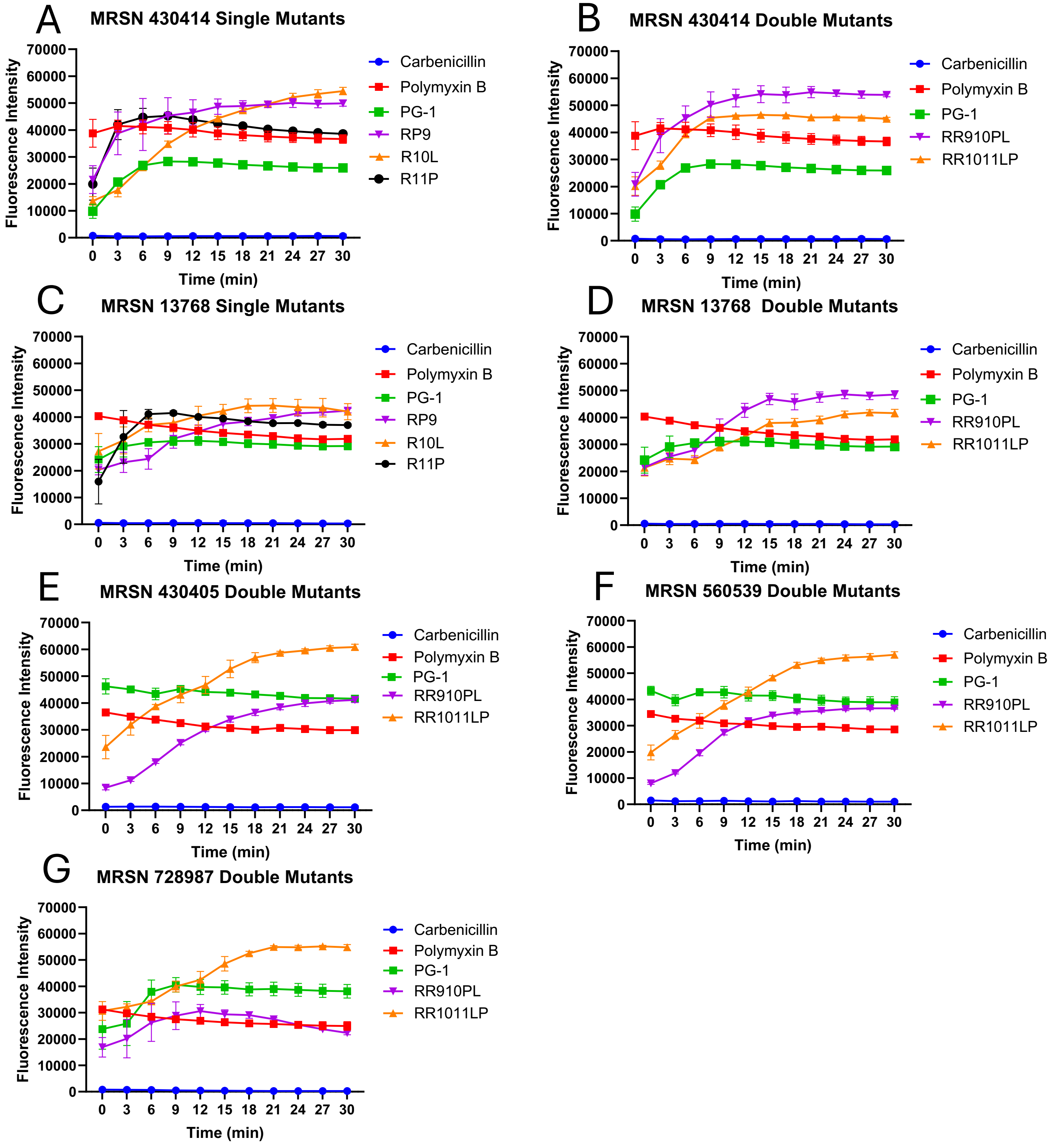


**Figure S4.** Outer Membrane Permeability with MRSN Clinical Isolates displaying elevated colistin resistance shows double mutant variant switch. Single mutant peptide variants evaluated against MRSN430414 (A) and MRSN 13768 (B). Double mutant peptide variants evaluated against MRSN430414 (C) MRSN13768 (D) MRSN 430405 (E) MRSN 560539 (F) and MRSN 728987. All assays were performed in triplicate with error shown as ±SEM.

**Figure S5 Dimer Center of Mass and Residue Contacts in GROMACs Simulations**

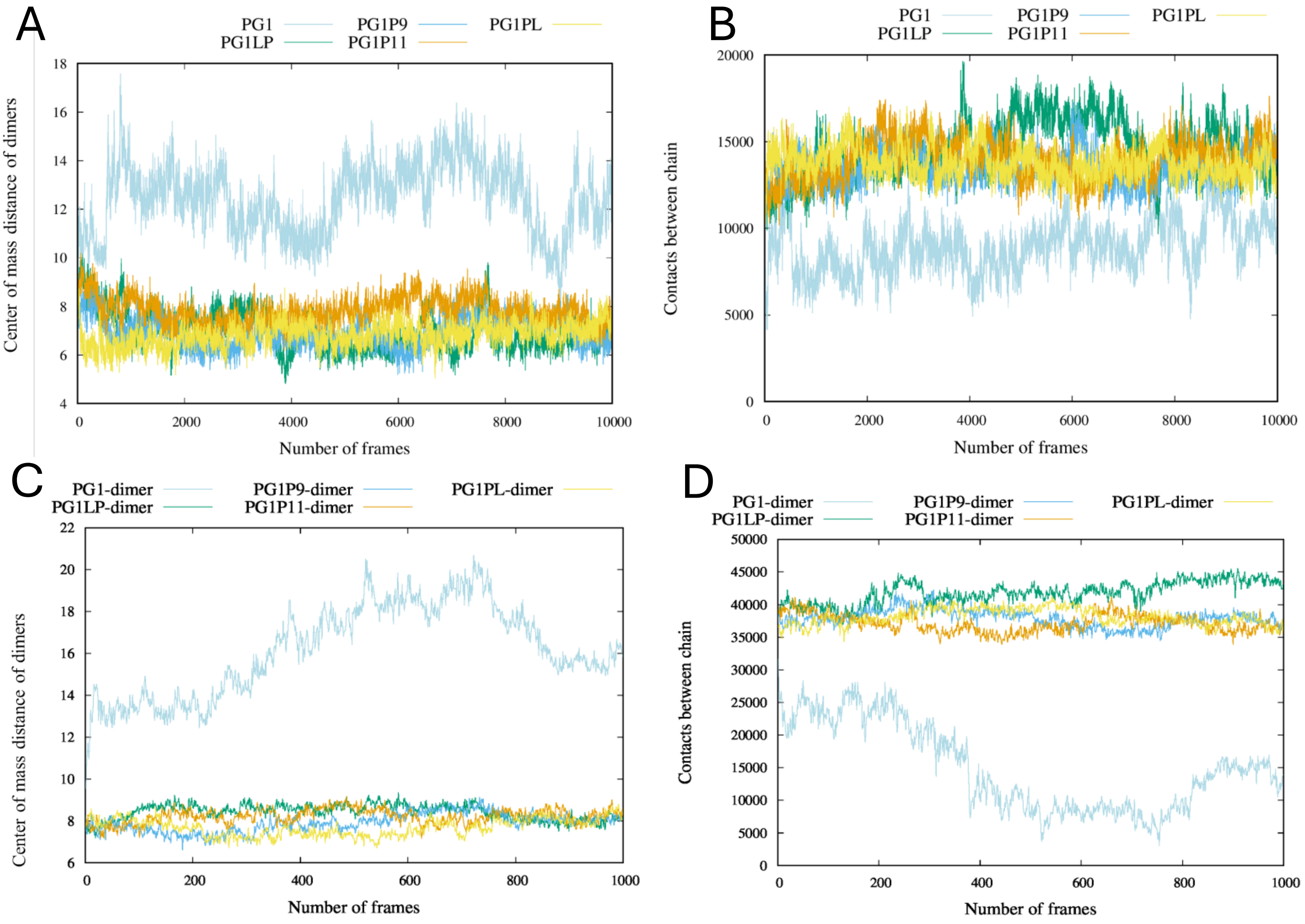


**Figure S5.** Center of Mass and Number of Contacts between Simulated dimers in salt box and in a general Gram-Negative inner membrane. Peptide variants were superimposed and simulated for 100ns and assessed for differences in distance between monomeric subunits over time in a salt box (0.15M NaCl at 310K) (A) or in a general Gram-Negative inner membrane (B). Peptide variants were also assessed for the number of interactions between each monomeric subunit in a salt box (0.15M NaCl at 310K) (C) or in a general Gram-Negative inner membrane (D)

### Tables

**Table S1. Bacterial Isolates**

| ***K. pneumoniae*** | **Characteristic** | **Isolation** | **Reference** |
|---|---|---|---|
| MKP103 | ST258; K107 | NIH clinical outbreak | **[47, 48]** |
| KPNIH1_10030-701::T30 | PhoP deficient | Transposon mutant | **[47]** |
| KPNIH1_07455-208::T30 | PagP deficient | Transposon mutant | **[47]** |
| KPNIH1_13050-101::T30 | PagL deficient | Transposon mutant | **[47]** |
| KPNIH1_24880-114::T30 | ArnT deficient | Transposon mutant | **[47]** |
| KPNIH1_08200-303::T30 | PmrB deficient | Transposon mutant | **[47]** |
| KPPR1S | K2 capsule serotype | ATCC 43816 , Rif$^r$ Str$^r$ | **[44]** |
| NTUH K2044 | K1 capsule serotype | Liver abscess | **[43]** |
| ATCC 13883 | Type strain; K3 | | **[46]** |
| ATCC 700603 | Clinical strain; K6 | Urine | **[45]** |
| MRSN 430414 | Colistin-resistant clinical isolate | Blood | **[50]** |
| MRSN 13768 | Colistin-resistant clinical isolate | Blood | **[50]** |
| MRSN 430405 | Colistin-resistant clinical isolate | Blood | **[50]** |
| MRSN 560539 | Colistin-resistant clinical isolate | Urine | **[50]** |
| MRSN 728987 | Colistin-resistant clinical isolate | Wound | **[50]** |
| *Burkholderia thailandensis* E264 | Drug-resistant soil bacterium | Rice Fields | **[65]** |
| *Acinetobacter baumannii* 5075 | Drug-resistant clinical isolate | Tibia/osteomyelitis | **[66]** |
| *Escherichia coli* W3110 | Type strain; K12 | | **[60], [67]** |
| *Escherichia coli* 25922 | Clinical Strain; O6 | | **[68]** |

**Table S2. Mixed Base Primer Sequence**

| | |
|---|---|
| Primer Sequence | 5' CTG CAG GTC GAC TTA (N1)(N2)(N3) (N4)(N2)(N2) (N4)(N1)(N2) R(N2)(N1) (N1)(N1)(N2) R(N2)(N1) (N3)(N1)(N1) (N2)(N2)(N4) (N3)(N2)(N3) (N1)(N2)(N3) R(N2)(N1) R(N4)(N1) R(N2)(N1) (N1)(N1)(N3) (N1)(N2)(N3) (N2)(N2)(N2) (N1)(N2)(N2) (N3)(N2)(N3) GGT TCC TCC GAT ACC CGC AG-3' |
| Reference for Nucleotide Conservation | N1 = (A 94%, T 2%, G 2%, C 2%)<br>N2 = (A 2%, T 94%, G 2%, C 2%)<br>N3 = (A 2%, T 2%, G 94%, C 2%)<br>N4 = (A 2%, T 2%, G 2%, C 94%) |

**Table S3. Barcodes used for Illumina sequencing**

| Tube # | Plasmid # (= template) | Illumina (Reverse) Primer # | Barcode |
|---|---|---|---|
| 1 | 1 | 7 | CAGATC |
| 2 | 2 | 8 | ACTTGA |
| 3 | 3 | 9 | GATCAG |
| 4 | 4 | 15 | ATGTCA |
| 5 | 5 | 18 | GTCCGC |
| 6 | 6 | 19 | GTGAAA |

**Table S4. Top 20 Inactive Peptide Sequences with Highest Log2Fold Change**

| baseMean[a] | log2FoldChange[b] | lfcSE[c] | Stat[d] | Pvalue[e] | Padj[f] | Amino Acid Sequence |
|---|---|---|---|---|---|---|
| 5.667618 | 3.129748492 | 0.96891825 | 3.2301471 | 0.001237 | 0.99998 | RGGRLNYCRRRLCVCVGL |
| 3.248165276 | 2.796891478 | 2.32963862 | 1.2005688 | 0.229919 | 0.99998 | RRGRLCYCPRRLCVCVGR* |
| 4.281980156 | 2.584835626 | 1.04488489 | 2.4737994 | 0.013368 | 0.99998 | RRARICYCPLRLRVCVGR* |
| 3.510769682 | 2.276195306 | 1.07017543 | 2.1269366 | 0.033425 | 0.99998 | GRGRLCYCRRRLCVCVVR* |
| 4.246091978 | 2.270721692 | 0.96593098 | 2.3508115 | 0.018733 | 0.99998 | RRGRICYCPLRVCVCVGR* |
| 2.550375501 | 2.167095837 | 1.2646099 | 1.7136477 | 0.086593 | 0.99998 | RRGRICYCPHRLCGCVGR* |
| 3.343462933 | 2.110018587 | 1.11478806 | 1.8927531 | 0.058391 | 0.99998 | RRGRIRYCPLRLCVCVGR* |
| 8.840698337 | 2.032328152 | 0.67399336 | 3.0153534 | 0.002567 | 0.99998 | RRGRICYCPLRLGVCVGR* |
| 4.570858954 | 1.995224781 | 0.92460115 | 2.1579302 | 0.030933 | 0.99998 | RRGRHRYCRRRLCVCVGR* |
| 2.708259667 | 1.932415503 | 1.19669858 | 1.6147888 | 0.106356 | 0.99998 | RGGRSCYCPRRLCVCVGR* |
| 2.660194258 | 1.898287824 | 1.21514001 | 1.5621968 | 0.118242 | 0.99998 | RRGRPCYCRRRLYVCVER* |
| 3.856757869 | 1.88890246 | 1.70201334 | 1.1098047 | 0.267083 | 0.99998 | RRRRICYCPLRCCVCVGR* |
| 3.374223489 | 1.869974867 | 1.04059044 | 1.7970325 | 0.07233 | 0.99998 | RGGRLCYCPLRYCVCVGR* |
| 1.949778362 | 1.792347371 | 1.44361499 | 1.2415688 | 0.214396 | 0.99998 | RRGRNCYCPLRFCVCVGR* |
| 3.8086216 | 1.775553179 | 0.98104551 | 1.8098581 | 0.070318 | 0.99998 | RGGRLCYCRRRLCVCVVR* |
| 4.542692666 | 1.739620454 | 0.89603069 | 1.9414742 | 0.052201 | 0.99998 | RRGRICYCPLRLCVCVGH* |
| 11.2423873 | 1.731554121 | 0.57047124 | 3.0353048 | 0.002403 | 0.99998 | RRGRIGYCPLRLRVCVGR* |
| 4.333506368 | 1.723282449 | 0.9072632 | 1.8994295 | 0.057508 | 0.99998 | RRGRMCYCPLRLGVCVGR* |
| 2.361088324 | 1.699166377 | 1.25229997 | 1.3568366 | 0.174833 | 0.99998 | RRGHIGYCPLRLCVCVGR* |
| 7.475621655 | 1.666379119 | 0.70853571 | 2.3518633 | 0.01868 | 0.99998 | RRGRICYCPLRLCVCVGR* |
| 5.272936824 | 1.597627819 | 0.83457013 | 1.9143123 | 0.05558 | 0.99998 | RRGRICYCPLRLGVCVGR* |
| 5.620389029 | 1.593919017 | 0.81607825 | 1.9531448 | 0.050802 | 0.99998 | RRGRTCYCPLRFCVCVGR* |
| 4.00507873 | 1.589101325 | 0.94705859 | 1.6779335 | 0.09336 | 0.99998 | RGGRLCYSRRRLRVCVGR |
| 5.339745145 | 1.583496038 | 0.81976488 | 1.9316466 | 0.053403 | 0.99998 | RRGRLCYCRRRLRVCVGR* |
| 9.807106054 | 1.579683216 | 0.65852601 | 2.3988167 | 0.016448 | 0.99998 | RRGRICYCPLRLRVCVGR* |

baseMean[a] is the averaged of the normalized counts across all samples

log2FoldChange[b] is the change in expression of a sequence relative to the uninduced population.

lfcSE[c] is the standard error value

Stat[d] is the Wald statistic (log2FolChange/ lfcSE)

Pvalue[e] is the overall significance of the model.

Padj[f] is the adjusted p value using the Benjamini and Hochberg method

**Table S5. Primer List**

| Oligo Name | Sequence |
|---|---|
| RpoDF | GATCTGATCACCGGTTTCGT |
| RpoDR | CTTCGTCGTCATCCATCTCTTC |
| PhoPF | GCCGGATGAAGACGGTTTAT |
| PhoPR | CGCTCAGCACTTCCACTTTA |
| pmmB67EHse5_2 | ggtcgtaaatcactgcataattcgtgtcgc |
| pmmB67EHseq3 | cgcagaagcggtctgataaaacagaatttg |
| PG-1 | ATAgtcgaCTTAACGTCCTACGCAAACACAGAAACGGCGACGGCAGTAACAAAGACGCCCACC GCGGGtTCCTCCGATACCcGCaGCtGGaGCC |
| PG1G2R | ctgcaggtcgacttaGCGTCCAACGCAAACACAAAAGCGACGACGGCAGTAACATAAGCGACCGC GACGggatccGATACCcGCaGCtGG |
| PG1L5I | ctgcaggtcgacttaACGACCAACGCACACACAGAAACGGCGACGACAGTAACAGATGCGACCC CCACGggatccGATACCcGCaGCtGG |
| PG1R9P | ctgcaggtcgacttaACGACCCACGCACACGCAAAAACGACGTGGACAGTAACACAGACGGCCTC CACGggatccGATACCcGCaGCtGG |
| PG1R10L | ctgcaggtcgacttaGCGACCAACGCATACACAGAAGCGTAAGCGACAATAACATAAGCGACCCC CACGggatccGATACCcGCaGCtGG |
| PG1F12L | ctgcaggtcgacttaGCGTCCCACACACACACACAAACGACGACGACAGTAACATAAACGACCGC CGCGggatccGATACCcGCaGCtGG |
| PG1RR910PL | ctgcaggtcgacttaACGGCCTACACACACGCAAAACCGAAGCGGGCAGTAACACAATCTTCCTC CGCGggatccGATACCcGCaGCtGG |

**Table S6. Plasmids used in this work**

| Plasmids name | Source |
|---|---|
| pMMB67EH_*lpp_ompA* | [59] |
| pMMB67EH _lpp_ompA_2xtether | [59] |
| pMMB67EH_*lpp_ompA_2x*tether_*protegrin 1* | [59] |
| pMMB67EH_*lpp_ompA_2 tether_R9P* | This Study |
| pMMB67EH_*lpp_ ompA_2xtether_R2G* | This Study |
| pMMB67EH_*lpp_ompA_2xtether_L5I* | This Study |
| pMMB67EH_*lpp_ompA_2xtether_R10L* | This Study |
| pMMB67EH_*lpp_ompA_2x tether_F12L* | This Study |
| pMMB67EH _*lpp_ompA_2xtether_RR910PL* | This Study |

**Table S7. Mult-drug-Resistant Organism Repository and Surveillance Network (MRSN) Colistin MICs**

| | Catalog Number | Part of World | Isolation point | Strain | VIR[a] | KL-type[b] | Colistin MIC $\mu$mol $L^{-1}$ /$\mu$g $mL^{-1}$ |
|---|---|---|---|---|---|---|---|
| 14 | NR-55517 | Europe | Blood | MRSN 13768 | 1 | 17 | 27.69/32 |
| 53 | NR-55556 | Middle East | Blood | MRSN 430414 | 1 | 15 | 27.69/32 |
| 52 | NR-55555 | Middle East | Blood | MRSN 430405 | 0 | 107 | 13.85/16 |
| 68 | NR-55571 | North America | Urine | MRSN 560539 | 0 | 36 | 13.85/16 |
| 91 | NR-55594 | Asia | Wound | MRSN 728987 | 0 | 122 | 13.85/16 |
| 41 | NR-55544 | Asia | Urine | MRSN 365679 | 3 | 51 | 6.92/8 |
| 42 | NR-55545 | Africa | Urine | MRSN 366562 | 1 | 64 | 6.92/8 |
| 43 | NR-55546 | Africa | Wound | MRSN 368001 | 0 | 2 | 6.92/8 |
| 44 | NR-55547 | Africa | Blood | MRSN 368320 | 1 | 52 | 6.92/8 |
| 45 | NR-55548 | Europe | Wound | MRSN 371351 | 0 | 57 | 6.92/8 |
| 46 | NR-55549 | North America | Urine | MRSN 374613 | 1 | 5 | 6.92/8 |
| 47 | NR-55550 | North America | Urine | MRSN 375436 | 2 | 3 | 6.92/8 |
| 54 | NR-55557 | North America | Urine | MRSN 450199 | 0 | 113 | 6.92/8 |
| 71 | NR-55574 | North America | Blood | MRSN 572640 | 2 | 2 | 6.92/8 |
| 99 | NR-55602 | Europe | Perianal | MRSN 752729 | 4 | 64 | 6.92/8 |
| 2 | NR-55505 | North America | Perianal | MRSN 4111 | 0 | 31 | 3.46/4 |
| 23 | NR-55526 | Asia | Wound | MRSN 19073 | 0 | 28 | 3.46/4 |
| 26 | NR-55529 | North America | Urine | MRSN 21352 | 0 | 22 | 3.46/4 |
| 33 | NR-55536 | South America | Urine | MRSN 27106 | 0 | 119 | 3.46/4 |
| 34 | NR-55537 | North America | Unknown | MRSN 27778 | 0 | 110 | 3.46/4 |
| 35 | NR-55538 | North America | Wound | MRSN 27989 | 0 | 3 | 3.46/4 |
| 36 | NR-55539 | North America | Respiratory | MRSN 28183 | 0 | 154 | 3.46/4 |
| 38 | NR-55541 | North America | Urine | MRSN 28880 | 1 | 27 | 3.46/4 |
| 51 | NR-55554 | North America | Urine | MRSN 414780 | 0 | 21 | 3.46/4 |

| | | | | | | | |
|---|---|---|---|---|---|---|---|
| 56 | NR-55559 | Asia | Wound | MRSN 479404 | 1 | 51 | 3.46/4 |
| 65 | NR-55568 | Europe | Urine | MRSN 526410 | 0 | 27 | 3.46/4 |
| 77 | NR-55580 | North America | Urine | MRSN 599975 | 1 | 2 | 3.46/4 |
| 92 | NR-55595 | North America | Blood | MRSN 730567 | 0 | 46 | 3.46/4 |
| 22 | NR-55525 | North America | Urine | MRSN 18411 | 0 | 14 | 1.73/2 |
| 25 | NR-55528 | North America | Urine | MRSN 21304 | 1 | 28 | 1.73/2 |
| 32 | NR-55535 | North America | Wound | MRSN 25947 | 0 | 30 | 1.73/2 |
| 40 | NR-55543 | North America | Urine | MRSN 28893 | 0 | 125 | 1.73/2 |
| 49 | NR-55552 | North America | Fluid | MRSN 401050 | 0 | 51 | 1.73/2 |
| 70 | NR-55573 | North America | Urine | MRSN 564304 | 0 | 62 | 1.73/2 |
| 86 | NR-55589 | Asia | Urine | MRSN 681054 | 1 | 15 | 1.73/2 |
| 94 | NR-55597 | North America | Urine | MRSN 736213 | 5 | 2 | 1.73/2 |
| 1 | NR-55504 | North America | Perianal | MRSN 1912 | 1 | 25 | 0.87/1 |
| 3 | NR-55506 | North America | Urine | MRSN 4759 | 0 | 38 | 0.87/1 |
| 5 | NR-55508 | North America | Urine | MRSN 5613 | 0 | 148 | 0.87/1 |
| 6 | NR-55509 | North America | Respiratory | MRSN 5741 | 2 | 30 | 0.87/1 |
| 7 | NR-55510 | North America | Wound | MRSN 5881 | 3 | 62 | 0.87/1 |
| 8 | NR-55511 | Europe | Wound | MRSN 6031 | 0 | 17 | 0.87/1 |
| 10 | NR-55513 | North America | Wound | MRSN 7076 | 0 | 25 | 0.87/1 |
| 11 | NR-55514 | Europe | Urine | MRSN 13726 | 0 | 23 | 0.87/1 |
| 12 | NR-55515 | Europe | Blood | MRSN 13748 | 1 | 10 | 0.87/1 |
| 13 | NR-55516 | Europe | Wound | MRSN 13761 | 0 | 3 | 0.87/1 |
| 16 | NR-55519 | North America | Urine | MRSN 15219 | 0 | 60 | 0.87/1 |
| 21 | NR-55524 | Asia | Urine | MRSN 16233 | 3 | 2 | 0.87/1 |
| 24 | NR-55527 | North America | Respiratory | MRSN 20522 | 1 | 24 | 0.87/1 |
| 27 | NR-55530 | South America | Respiratory | MRSN 22232 | 1 | 151 | 0.87/1 |

| | | | | | | | |
|---|---|---|---|---|---|---|---|
| 28 | NR-55531 | South America | Respiratory | MRSN 22265 | 0 | 102 | 0.87/1 |
| 29 | NR-55532 | North America | Urine | MRSN 25107 | 0 | 30 | 0.87/1 |
| 30 | NR-55533 | North America | Urine | MRSN 25112 | 0 | 52 | 0.87/1 |
| 31 | NR-55534 | North America | Urine | MRSN 25616 | 0 | 19 | 0.87/1 |
| 48 | NR-55551 | North America | Urine | MRSN 380979 | 0 | 25 | 0.87/1 |
| 55 | NR-55558 | North America | Urine | MRSN 468268 | 0 | 58 | 0.87/1 |
| 57 | NR-55560 | North America | Urine | MRSN 499958 | 0 | 10 | 0.87/1 |
| 58 | NR-55561 | Middle East | Unknown | MRSN 511348 | 0 | 2 | 0.87/1 |
| 59 | NR-55562 | Middle East | Unknown | MRSN 513382 | 0 | 30 | 0.87/1 |
| 60 | NR-55563 | Middle East | Unknown | MRSN 515247 | 2 | 1 | 0.87/1 |
| 62 | NR-55565 | Middle East | Unknown | MRSN 516635 | 0 | 30 | 0.87/1 |
| 64 | NR-55567 | Middle East | Unknown | MRSN 518712 | 0 | 61 | 0.87/1 |
| 66 | NR-55569 | North America | Blood | MRSN 539414 | 0 | 9 | 0.87/1 |
| 67 | NR-55570 | North America | Urine | MRSN 546733 | 0 | 46 | 0.87/1 |
| 69 | NR-55572 | North America | Urine | MRSN 562722 | 2 | 116 | 0.87/1 |
| 72 | NR-55575 | Asia | Urine | MRSN 581745 | 0 | 35 | 0.87/1 |
| 78 | NR-55581 | North America | Urine | MRSN 607210 | 0 | 51 | 0.87/1 |
| 79 | NR-55582 | Africa | Environmental | MRSN 613682 | 1 | 112 | 0.87/1 |
| 80 | NR-55583 | Africa | Environmental | MRSN 614201 | 0 | 14 | 0.87/1 |
| 81 | NR-55584 | Europe | Blood | MRSN 669448 | 1 | 102 | 0.87/1 |
| 83 | NR-55586 | North America | Urine | MRSN 672476 | 0 | 39 | 0.87/1 |
| 84 | NR-55587 | North America | Urine | MRSN 676980 | 0 | 30 | 0.87/1 |
| 85 | NR-55588 | North America | Urine | MRSN 680172 | 0 | 46 | 0.87/1 |
| 87 | NR-55590 | North America | Urine | MRSN 699478 | 0 | 14 | 0.87/1 |
| 88 | NR-55591 | North America | Urine | MRSN 699654 | 0 | 65 | 0.87/1 |
| 90 | NR-55593 | Africa | Wound | MRSN 702325 | 1 | 107 | 0.87/1 |

| | | | | | | | |
|---|---|---|---|---|---|---|---|
| 93 | NR-55596 | Asia | Urine | MRSN 731029 | 3 | 15 | 0.87/1 |
| 95 | NR-55598 | Europe | Respiratory | MRSN 740795 | 1 | 2 | 0.87/1 |
| 97 | NR-55600 | North America | Urine | MRSN 750877 | 0 | 62 | 0.87/1 |
| 98 | NR-55601 | North America | Urine | MRSN 750999 | 0 | 107 | 0.87/1 |
| 100 | NR-55603 | North America | Blood | MRSN 761403 | 0 | 107 | 0.87/1 |
| 4 | NR-55507 | North America | Wound | MRSN 4815 | 0 | 38 | 0.43/0.5 |
| 9 | NR-55512 | North America | Urine | MRSN 6778 | 0 | 3 | 0.43/0.5 |
| 15 | NR-55518 | North America | Urine | MRSN 14444 | 0 | 142 | 0.43/0.5 |
| 17 | NR-55520 | North America | Urine | MRSN 15687 | 0 | 62 | 0.43/0.5 |
| 18 | NR-55521 | Europe | Perianal | MRSN 15882 | 0 | 113 | 0.43/0.5 |
| 19 | NR-55522 | North America | Wound | MRSN 15937 | 0 | 42 | 0.43/0.5 |
| 20 | NR-55523 | North America | Urine | MRSN 16008 | 1 | 63 | 0.43/0.5 |
| 37 | NR-55540 | North America | Wound | MRSN 28866 | 0 | 146 | 0.43/0.5 |
| 39 | NR-55542 | North America | Urine | MRSN 28887 | 0 | 15 | 0.43/0.5 |
| 50 | NR-55553 | North America | Urine | MRSN 410359 | 0 | 57 | 0.43/0.5 |
| 61 | NR-55564 | Middle East | Unknown | MRSN 515432 | 4 | 43 | 0.43/0.5 |
| 63 | NR-55566 | Middle East | Unknown | MRSN 517281 | 0 | 64 | 0.43/0.5 |
| 73 | NR-55576 | Asia | Respiratory | MRSN 582610 | 5 | 20 | 0.43/0.5 |
| 74 | NR-55577 | Europe | Perianal | MRSN 583114 | 0 | 28 | 0.43/0.5 |
| 75 | NR-55578 | North America | Urine | MRSN 583141 | 0 | 13 | 0.43/0.5 |
| 76 | NR-55579 | North America | Wound | MRSN 591344 | 0 | 61 | 0.43/0.5 |
| 82 | NR-55585 | Europe | Respiratory | MRSN 669510 | 0 | 62 | 0.43/0.5 |
| 89 | NR-55592 | Africa | Wound | MRSN 702261 | 1 | 54 | 0.43/0.5 |
| 96 | NR-55599 | North America | Blood | MRSN 742743 | 0 | 128 | 0.43/0.5 |

VIR[a] is the virulence score given to each isolate with 5 being extremely variant

KL-type[b] is the capsule type of each isolate

**Table S8. Minimum Inhibitory Concentrations Other Gram-Negative Bacteria (μmol $L^{-1}$/ μg $mL^{-1}$)**

| Bacterial Strain | Polymyxin B | Protegrin-1 | R9P | R11P | R9P,R10L | R10L,R11P |
|---|---|---|---|---|---|---|
| *B. thailandensis E264* | >49.17/>64 | >29.62/>64 | >30.45/>64 | >30.45/>64 | >31.09/>64 | >31.09/>64 |
| *A. baumannii 5075* | 0.77/1 | 0.46/1 | 0.95/2 | 0.48/1 | 7.77/16 | 0.97/2 |
| *E. coli W3110* | 0.77/1 | 0.46/1 | 0.95/2 | 0.95/2 | 3.89/8 | 1.94/4 |
| *E. coli 25922* | 0.77/1 | 0.93/2 | 3.81/9 | 0.48/1 | 3.89/8 | 0.97/2 |
